# Intrinsically disordered regions influence protein motif usage in *Drosophila* Ubx and AbdA Hox proteins

**DOI:** 10.64898/2026.10.09.757964

**Authors:** Corinne Maurel-Zaffran, Ilan Edelstein, Laurianne Michel, Meiggie Macchi, Nathalie Caruso, Andrew J Saurin, Yaakov Levy, Yacine Graba

**Affiliations:** Aix-Marseille Univ, CNRS, Developmental Biology Institute of Marseille (IBDM), IBDM-UMR 7288, Case 907, Parc Scientifique de Luminy, Marseille Cedex 09 13288, France; Dept. of Chemical and Structural Biology, Weizmann Institute of Science, Rehovot, Israel

**Keywords:** Hox, IDR, development, gene regulation, Drosophila

## Abstract

Hox proteins are evolutionary conserved transcription factors guiding morphogenesis in all bilaterian animals. Conservation mainly concerns the homeodomain (HD), a helix-turn-helix DNA binding motif and flanking sequences close by. Besides, sequences are poorly conserved and disordered. Here we investigate how poorly conserved Intrinsically Disordered Regions (IDRs) influence Hox protein activity, focusing on a closely related *Drosophila* Hox pair and using a wide range of in vivo function. Studying *Drosophila* Ultrabithorax (Ubx) and AbdominalA (AbdA) proteins, we find that conserved protein motifs close to the HD are differently used in Ubx and AbdA. Coarse-grained molecular dynamics simulations identify a large set of intra-molecular interactions, notably involving poorly conserved disordered regions upstream and downstream the HD. Swapping these regions in Ubx and AbdA alters protein functions and switches motif dependency in some biological situations. This suggests that IDRs, predicted to mediate large and dynamic sets of intramolecular interactions, influence protein motif usage in a context dependent manner.

## Introduction

Hox genes encode key transcription factors controlling morphological diversification in all bilaterians (Krumlauf, 1994; Rezsohazy et al., 2015). They have been the focus of extensive studies, to apprehend the breath of their developmental and evolutionary functions, but also to grasp how regulatory information is encoded within protein sequence. Hox proteins share a highly conserved 60 amino acids helix-turn-helix motif, the homeodomain (HD) that serves DNA binding, recognizing a low complexity DNA motif, TAAT, which contrasts with their in vivo functional specificity. Most Hox proteins also share a short protein interaction motif, the Hexapeptide (HX), lying close to and upstream of the HD. While also serving other functions (Baeza et al., 2015; Merabet et al., 2011; Prince et al., 2008), the best characterized function of the HX is to contact PBC class cofactors, forming a protein complex with improved DNA binding selectivity (Bobola and Merabet, 2016; Mann and Chan, 1996; Merabet and Mann, 2016; Merabet s., 2009).

The HD and HX motif represent only a small portion of Hox protein sequence, which in size, in *Drosophila* for example, ranges from 330 to 782 amino acids. How disordered protein regions impact on Hox protein function has been addressed only in a few studies. Sequences analyses combined to disorder prediction very early identified that most of Hox proteins sequences appear disordered (Liu et al., 2008). Predictions for SLiMs, Short Linear Motifs located within Intrinsically Disordered Regions (IDRs), further revealed a widespread distribution of SLiMs along Hox protein sequences, and identified that they define paralog specific signatures, suggesting a contribution to Hox protein specificity (Rinaldi et al., 2018). Work based on in vitro, yeast two hybrid and in embryo Bi-fluorescent complementation assays (BiFC) showed that the disordered protein regions widely contribute to define Hox interactome (Baeza et al., 2015; Dard et al., 2018). Among all Hox proteins, IDRs have been best characterized in the case of the *Drosophila* Ultrabithorax (Ubx) protein. A combination of biophysical, biochemical, biocomputing and yeast interaction assays indicated that disordered regions establish contact with protein partners (Hsiao et al., 2014). These interactions rely on the simultaneous use of multiple Ubx IDRs, that promote contacts towards proteins that share some topological similarities. Similar approaches including in vitro DNA binding studies, always on Ubx, and probing IDRs in isolation or combinations, further establish a complex role of IDRs on the control of Ubx DNA binding (Liu et al., 2008). These data argue for a widespread contribution of disordered protein sequences to Hox protein function.

To explore further the role of IDRs in Hox proteins, we focused on a pair of proteins whose sequence mainly diverges within disordered protein regions. *Drosophila* Ubx and Abdominal-A (AbdA) are central class Hox proteins with the highest sequence similarities within the *Drosophila* Hox complement (Hueber et al., 2010). The HDs of Ubx and AbdA differs in only 4 residues out of 60 (Chauvet et al., 2000). Ubx and AbdA also share a short peptide immediately adjacent and downstream to the HD, only present in Ubx and AbdA, termed UbdA ((Balavoine et al., 2002); UA, a contraction between Ubx and AbdA, Fig. 1A, B). This domain emerged at the basis of protostomes, and became strongly conserved in pan-arthropods (arthropods and onycophorans, Fig. 1B). No such domains are shared in any other *Drosophila* Hox proteins, defining Ubx/AbdA as a unique phylogenetic pair. Structural and functional studies of the UA motif indicate that it provides weak and plastic contacts towards the Exd cofactor (Foos, 2015; Ortiz-Lombardia et al., 2017). As for the HX motif, the function of the UA motif is not restricted to contact Exd, as many Exd-independent function of Ubx and AbdA requires the UA motif (Baeza et al., 2015; Merabet et al., 2011; Prince et al., 2008). As other Hox proteins, Ubx and AbdA proteins are mostly disordered outside the HD and its flanking sequences (Fig. 1C).

**Figure 1.**
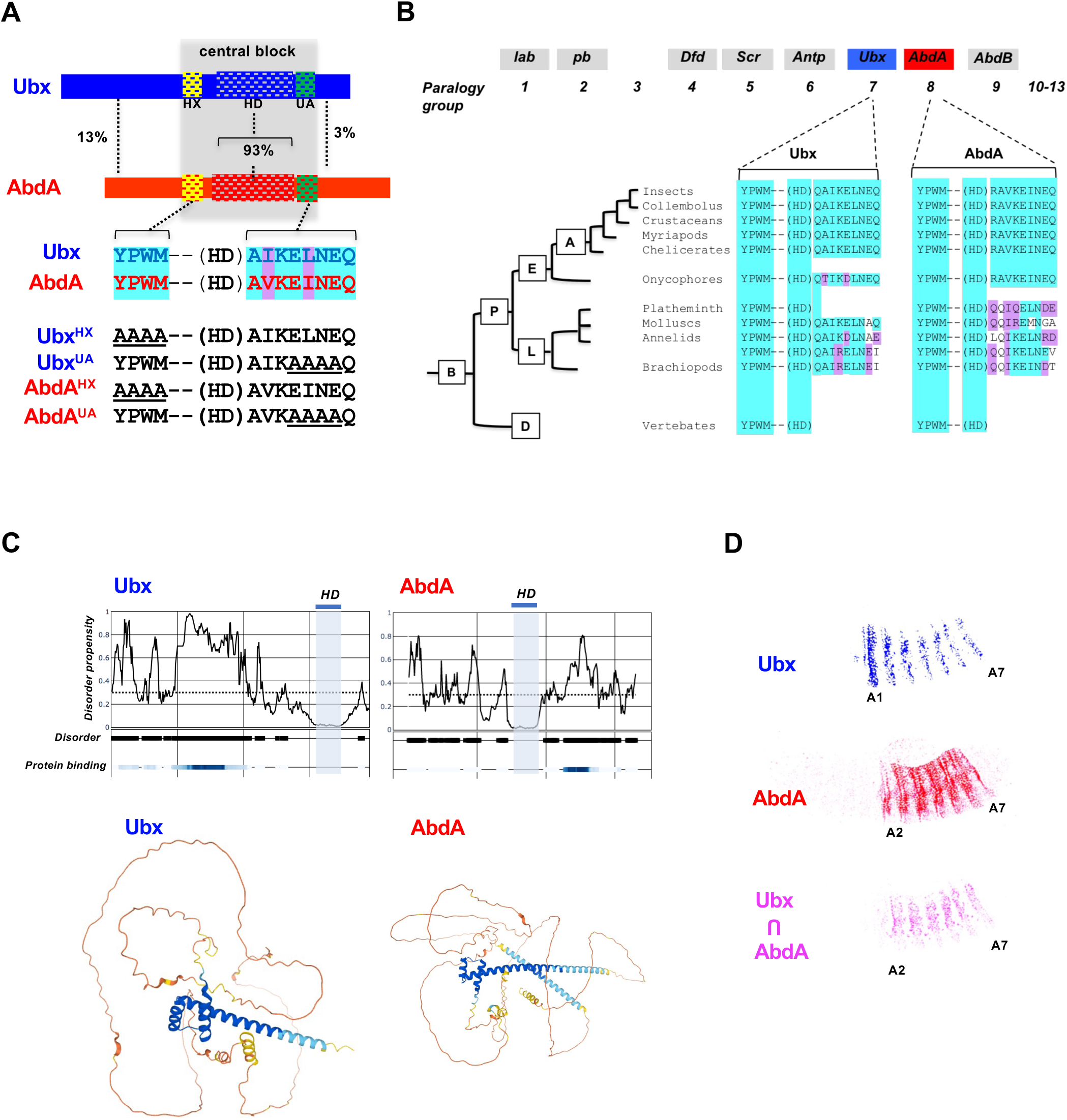
Ubx and AbdA are closely related Hox proteins. **(A)** Ubx/AbdA sequence conservation. Conservation applies to the HD that displays 93% of sequence identity and the HX and UA motifs, defining a conserved HD centered central block. Sequence identity is highlighted in cyan and similarities in purple. N and C-terminal sequences are poorly conserved. Sequences and mutations of the HX and UA motifs used in this study are indicated. **(B)** The Ubx/AbdA HD central block is evolutionarily highly conserved in protostomes paralog group Hox 7-8. A, B, D, E, L and P stand respectively for arthropods, bilaterians, deuterostomes, ecdysozoa, lophotrocozoa and protostomes. Two sequences from annelids are shown to illustrate the identified sequence diversity. **C)** Ubx and AbdA disorder prediction using the flDPnn tool (Hu et al., 2021) that couple disorder and protein binding prediction (upper panels), and alpha fold 3D structure predictions (lower panels). **(D)** Ubx and AbdA are expressed in widely overlapping territories. Embryo were stained for Ubx (blue) and AbdA (red), and images to illustrate overlapping expression patterns (pink) were obtained as described in the method section. Overlapping expression territories (pink) were obtained by image superimposition (Fig. S12).

In addition to concentrate protein sequence differences within IDRs, the Ubx /AbdA pair displays a second feature in being expressed in a similar embryonic pattern, covering segments T3 to A7 for Ubx (Akam and Martinez-Arias, 1985; White and Wilcox, 1985) and A2 to A7 for AbdA (Karch et al., 1990; Macias et al., 1990), thus overlapping in most abdominal segments (segments A2 to A7, Fig. 1D). These largely overlapping expression patterns allow the sharing of developmental functions key for patterning the epidermis, the central nervous system, the somatic muscles and the heart. This unique situation allows for probing experimentally whether divergence in IDRs impacts on Ubx and AbdA activity, within the frame of the same developmental functions, minimizing/alleviating the influence that distinct cellular context may provide. Besides common function, Ubx and AbdA bear specific functions that also relies on the integrity of the HX and/or UA motifs. This is illustrated by their activity in the visceral mesoderm, where Ubx expressed in the PS7 activates *decapentaplegic* (*dpp*) while AbdA expressed in PS8-12 represses *dpp* and activate *wingless* (Merabet et al., 2011; Merabet et al., 2007; Reuter et al., 1990).

In this study we perform a quantitative comparison of motif dependence across multiple Ubx and AbdA in vivo functions. Results highlight distinct motif requirements for Ubx and AbdA protein activities, which can be influenced by IDRs. We further use in silico molecular modeling to grasp intrinsic protein conformational properties, suggestive of how IDRs may influence motif dependency.

## Results

### Ubx and AbdA make different use of the HX and UA motifs

We explored four in vivo Ubx and AbdA shared functions that are known or likely Exd-independent (see Table S1). The choice was motivated by our strategy to combine in vivo data with Ubx and AbdA molecular modeling simulations performed in the absence of Exd. We expressed Hox proteins (WT, HX or UA mutants) in the anterior regions where endogenous Ubx or AbdA proteins are absent, avoiding functional interference, using the binary UAS-Gal4 system (Brand and Perrimon, 1993). Conditions allowing for comparable expression levels (+/-15%) were defined (see methods).

We started to investigate protein sequence requirements for denticle belt patterning. The *Drosophila* segmental organization is evidenced by denticles belts, that form ventrally within the anterior region of each segment, and that witnesses segment identity promoted by Hox proteins. In the first thoracic segment (T1) only, a group of smaller denticles (approximately 180, referred to as the beard), forms posterior to the main denticle belt (Fig. 2A). This is suppressed in the abdomen by Ubx and AbdA (Andrew et al., 1994), and arm-Gal4 driven anterior ectopic expression of Ubx or AbdA result in beard suppression within T1 (Fig. 2A) (Chan and Mann, 1993; Chauvet et al., 2000; Gibson et al., 1990). A quantitative assessment of the beard suppressive activity was obtained by counting the number of beard denticles following expression of WT or motif mutated proteins (see Table S2). Under our experimental conditions, the expression of the WT Ubx resulted in the total loss of beard denticles (scored as 100% of repressive activity) and that of AbdA resulted in an average of 6.7 remaining beard denticles, scored as 96% of repressive activity ((180-6.7)/180x)100)). Mutation of the HX or UA motifs in AbdA only very weakly affects the beard suppressive activity (average of 2.7 remaining denticles and 99% of activity for AbdA^HX^ and average of 17.1 remaining denticles and 90% for AbdA^UA^), while in the case of Ubx it reduces the suppressive activity to 41% (average of 105.8 remaining beard denticles) for the HX mutation, and 50% for the UA mutation (average of 89.8 remaining beard denticles) (Fig. 2A’). As the aim of the study is to assess motif contribution to protein function, we deduced motif contribution defined as the difference of protein activity in the wild type and motif-mutated protein (Table S2). When the mutated Hox activity equals that of the wild type protein, motif contribution equals zero; when the mutated Hox activity is weaker than that of the wild type protein, then motif contribution increases in proportion with the loss of mutated protein activity. Applied to beard suppression, HX and UA motif contribution to AbdA activity is close to zero, while it reaches 59% and 50% respectively for Ubx (Fig. 2A”). We conclude that while the HX and UA motifs are required for beard suppression by Ubx, this is not the case for AbdA.

**Figure 2.**
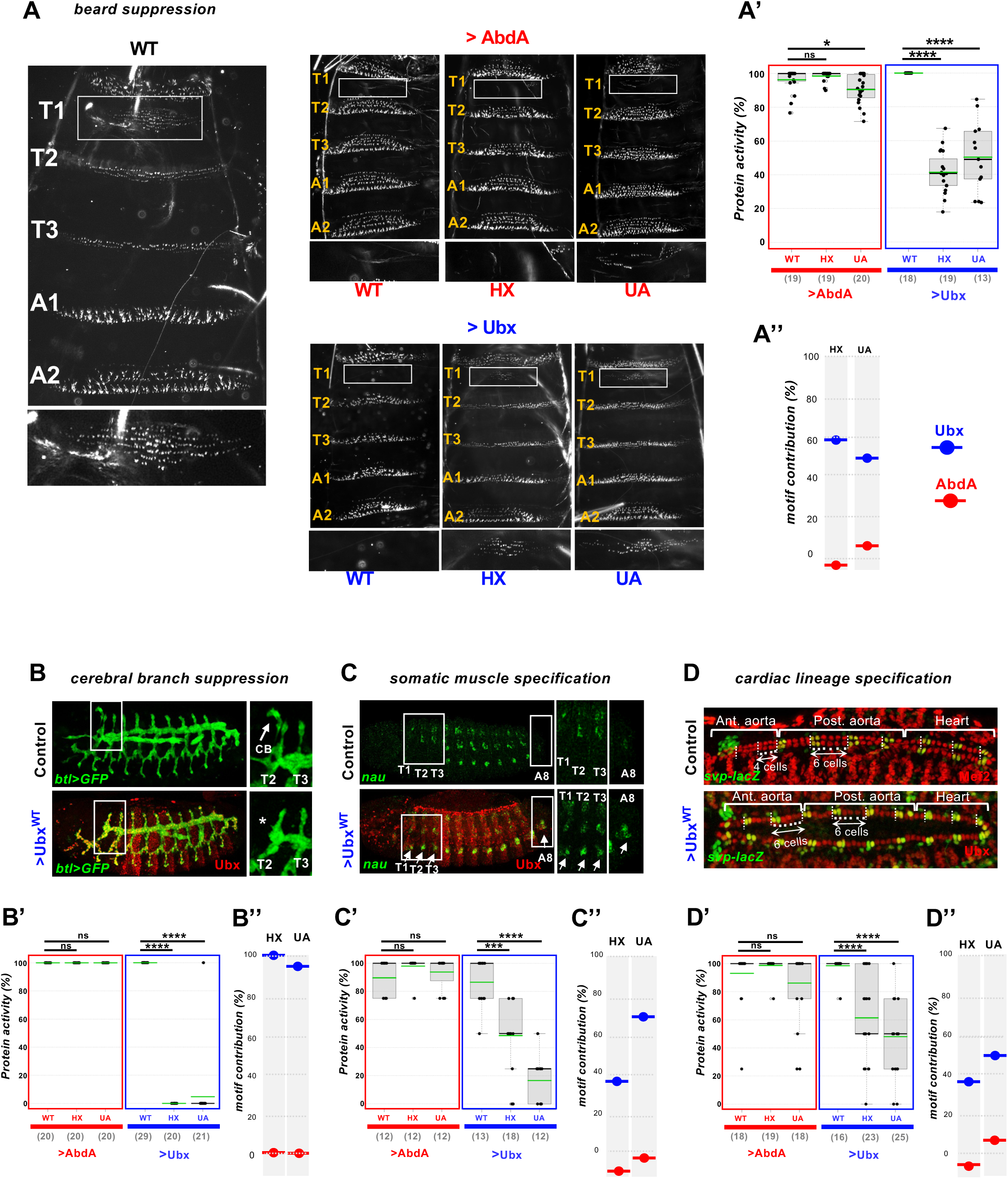
Shared Ubx and AbdA developmental functions rely on distinct HX and UA motif usage. (**A, A’, A”) Beard suppression**. **(A)** The beard, a group of around 180 small denticles, framed in a white rectangle, forms exclusively in the first thoracic segment T1 (A, left panels). A magnified view of the beard is shown below (A, left panel). *arm-Gal4* driven ectodermal expression of Ubx or AbdA suppress beard formation (A right panels). Yellow lettering indicates position of segments as referred to WT. Suppression is impaired by HX or UA mutations. **(A’)** The number of beard denticles in T1 were scored to define proteins activities in beard suppression **(A’’)**. Motif contribution was inferred from protein activity (HoxWT activity - HoxMut activity, Table S2) **(B, B’, B”) Cerebral branch repression. (B)** The cerebral branch exclusively forms dorsally in T2 (arrow). Ectopic expression of Ubx (or AbdA, see Fig. S1 and Table S2) suppress the formation of the cerebral branch (*). **(B’)** Presence or absenceof the tracheal branch in T2 was scored to define proteins activities. **(B”)** Inferred motif contributions (HoxWT activity - HoxMut activity, Table S2). **(C, C’, C”) Somatic muscle specification**. **(C)** Thoracic segments T1-T3 and Abdominal segment A8 lack a ventral somatic muscle cluster. Ectopic expression of Ubx or AbdA (see Fig. S2 and Table S2) most often induces ventral clusters in T1, T2, T3 and A8 (arrows). (**C’)** The number of induced ventral somatic muscle clusters (1 to 4) was scored to define protein activities. **(C”)** Inferred motif contributions (HoxWT activity - HoxMut activity, Table S2). **(D, D’, D”) Cardiac lineage specification. (D)** Hemi-segments (vertical white dashed lines) in the anterior aorta comprise 4 cells, while posterior aorta hemi-segments have 6. Ectopic expression of Ubx or AbdA (see Fig. S3 and Table S2) most often induces a 6 cell lineage in the anterior aorta. **(D’)** The number of cardiac cells in the two posterior most anterior aorta hemi-segments (8-12) were scored to define protein activities. **(D”)** Inferred motif contributions (HoxWT activity - HoxMut activity, Table S2). Numbers in brackets on graphs indicate the sample size (N) of each experiment. Statistical significance using Fisher or Mann-Withney tests (see methods) are **** for p<0.0001, *** for 0.0001<p<0.001, ** for 0.001<p<0.01, * for 0.01<p<0.05 and ns (not significant) for p>0.05. Mean values are in green.

To probe further if distinct motif usage is also at play for other shared Ubx and AbdA developmental functions, we applied a similar quantitative approach (see methods section) to three additional Ubx/AbdA shared functions. The first concerns the shaping of the larval respiratory system, the trachea. The cerebral branch is a small dorsally located branch of the tracheal network that exclusively forms in the second thoracic segment (T2), owing to posterior suppression by bithorax complex encoded proteins Ubx and AbdA, as well as AbdB (Chiang et al., 1995). The *breathless* (*btl*) Gal4 driver was used both as a tracheal reporter and as a driver for Hox expression. We scored either the presence or absence of tracheal branch in T2. In our experimental conditions (see methods), the expression of the WT Ubx or AbdA proteins resulted in all cases in the loss of cerebral branch (Fig. 2B and Fig. S1). Scores of cerebral branch repressive activity was defined by the frequency of cerebral branch repression. Both WT Ubx and AbdA thus were assigned with 100% tracheal repressive activity. Suppression of the cerebral branch by AbdA requires neither the integrity of the HX nor that of the UA motif (Fig. 2B’ and Fig. S1, (Merabet et al., 2011)). In contrast, in Ubx, mutation of the HX motif results in the total loss of cerebral branch suppression, and mutation in the UA motif in the large majority of cases also results in the loss of cerebral branch suppression, with very occasional cerebral branches observed following UA mutation (20 embryo of 21, Fig. 2B’ and Fig. S1). We concluded that the HX and UA motif do not contribute to AbdA-mediated cerebral branch repression, while they strongly do (100% for HX and 95% for UA) for Ubx (Fig. 2B’’ and Fig. S1).

The abdominal somatic musculature pattern is characterized by muscles located ventrally in the abdominal segments A1-A7 (Fig. 2C). These muscles are absent in the thorax and the abdominal segment A8. This is visualized by the expression of *nautilus* (*nau*). The A1-A7 abdominal specific pattern results from Ubx and AbdA activity (Michelson, 1994). Accordingly, expression of Ubx or AbdA using the mesodermal ubiquitous driver (*24B-Gal4*) results in ectopic ventral clusters of *nau* expressing cells anteriorly in all thoracic segments (T1-3) as well as posteriorly in abdominal segment A8. The quantitative assessment of abdominal type somatic muscle inducing potential was assessed by scoring the number of segment where the muscle pattern is transformed: A full transformation (protein activity equals 100%) is when a ventral muscle cluster is present in 4 segments, T1-T3 and A8; partial transformation, with 1 to 3 extra ventral muscle clusters were assigned protein activities of 75% to 25%; no transformation was assigned a score of 0%. In our experimental conditions (see methods), the expression of the WT Ubx or AbdA proteins resulted in transformation of at least 2, and most frequently 4 segments, with a mean of protein activity close to 90% in both cases (Fig. 2C’ and Fig. S2). Mutation of the HX or UA motifs in AbdA does not affect its capacity to promote ventral muscle clusters in T1-3 and A8 (Fig. 2C and Fig. S2; (Merabet et al., 2011). The situation is distinct for Ubx: HX mutation results in a protein that only promotes the appearance of ventral muscle in 2 or 3 additional segments, while UA mutation has a much stronger effect, rarely imposing ventral muscle clusters, at best only in a single segment. We concluded that while both the HX and UA motifs contribute to Ubx activity, they are dispensable for AbdA activity (Fig. 2C”).

The embryonic heart is patterned into anterior and posterior aorta and at its posterior end the heart proper. Segments in the posterior aorta and heart are made of six pairs of cells, instead of four in the anterior aorta (Fig. 2D). This difference is imposed by AbdA and Ubx (Perrin et al., 2004). Heart cell lineage is followed by *seven up* (*svp*) expression that allows delimiting segments. We counted the number of cardiac cells in the two posterior most segments of the anterior aorta on each side of the embryo (hemi embryo). A full transformation (protein activity equals 100%) was assigned when 12 cells are present (Fig. 2D’). Partial transformation, ranging from 11 to 9 cells were assigned protein activities of 75% to 25%, and lack of transformation (8 cells) protein activity of 0%. In our experimental conditions (see methods), mesodermal ubiquitous expression (*24B-Gal4*) of wild type Ubx or AbdA expression most often induces a strong transformation (respectively 98% and 93% of protein activity, Fig. 2D’ and Fig. S3). Mutation of the HX or UA motifs in AbdA does not affect the capacity of the protein to induce a six-cell lineage, with protein activities averaging 99% and 86%. In contrast, mutation of these motifs results in a partial loss of six cell lineage promoting capacity, with Ubx^HX^ and Ubx^UA^ protein activities declining to 61 and 48% (Fig. 2D’). We concluded that the HX and UA motif contribution to AbdA activity was close to zero, while their contribution to Ubx activity reaches 37% and 50% respectively (Fig. 2D”).

Overall, these data highlight that the HX and UA motifs are essential or strongly contribute to full Ubx activity, while they generally do not or only marginally contribute to AbdA activity. The core of the HX motif in Ubx and AbdA are identical, the core of the UA motif varies in two positions. Although the changes are conservative (I->V and L>I), the general lack/weak contribution of the UA motif may result from the motif being generally non-functional in AbdA, as a result of the two amino acid changes. We tested the functionality of the AbdA UA motif by probing if and to what extent it can restore the loss of activity induced by its mutation in Ubx. We modified these two positions within Ubx (as well as the 2 UA flanking amino acids to favor a genuine motif context) so that the engineered protein harbors an AbdA like UA motif (Ubx^UA(AbdA)^). The activity of Ubx^UA(AbdA)^ in the functional readouts previously used was determined (Fig. S4). Results show that the AbdA UA motif restores A1-A7 somatic muscle pattern induction (17 to 56% of activity) and cardiac six lineage commitment induction (48 to 93% of activity), indicating that the AbdA UA motif is not generally non-functional. The lack of functional restoration for beard suppressing activity and cerebral branch repression further indicates that in these biological situations, the precise identity of the UA motif matters.

We concluded that HX and UA motifs are differentially used in Ubx and AbdA, with a general requirement for Ubx and a general dispensability for AbdA. Importantly, these differences are not the consequence of Ubx and AbdA acting in different cellular context, as the impact of motif mutations is assessed for shared developmental functions consecutive to anterior ectopic expression within the same cellular fields.

### Molecular modeling predicts a dense network of intra-molecular interaction

The main sequence divergence between Ubx and AbdA that may be at the origin of HX and UA differential use lies outside the HD central block, with 13% and 3% of sequence conservation for the N-ter and C-ter sequences (Fig. 1A). These sequences are mainly composed of disordered protein regions, with Ubx notably displaying a large IDRs within the N-ter not found in AbdA (Fig. 1C).

To study the N and C-ter sequences that lack structural information, we performed in silico modeling using coarse-grained molecular dynamics simulations (Shea et al., 2021). Such simulations define preferential conformation ensembles and contact trajectories suggestive, without establishing causality, of dynamic protein features that may account for protein activities (Fuxreiter, 2018; Wright and Dyson, 2015). Simulations were done on full length Ubx and AbdA, with the aim of identifying intrinsic cues on which higher order information may further act. In our model, the homeodomain remains folded throughout the simulations and the IDRs are flexible and can participate in intramolecular interactions with both disordered regions or the homeodomain. The model includes hydrophobic interactions and electrostatic interactions between charged residues. This model, as well as other models with a similar approach were successfully applied to Intrinsically Disordered Proteins (IDPs) to predict protein compaction assessed by the radius of gyration, representing the radial distance of a sphere within which the protein fits (Dannenhoffer-Lafage and Best, 2021; Hazra et al., 2023; Joseph et al., 2021; Regy et al., 2021; Tesei et al., 2021). The Ubx and AbdA proteins were subdivided in three main regions, N-ter, C-ter and central block delimited by the HX and UA domains (Fig. 3A). Contact maps reporting the probability of contact between two residues were established with darker regions representing a higher probability of contact, indicative of a more compact conformation (Fig. 3B(I, II)). The contact map of AbdA appears qualitatively "darker" than that of Ubx, suggesting that AbdA adopts a more compact conformation compared to Ubx as illustrated by selected conformations in Fig. 3C(I, II). This conclusion is further supported by the global radius of gyration, Rg, of the proteins (Fig. 3D; WT) and the Rg of the N-ter IDR (Fig. 3E; WT)) of Ubx and AbdA, where AbdA appears more compact than Ubx. The C-ter IDRs do not follow this trend, but the very short C-ter in Ubx make the comparison less relevant. The Ubx and AbdA contact maps exhibit different and complex patterns. Within the N-ter sequences, Ubx displays a small number of regions (3 main ones) where the probability of contact within each region is high (Fig. 3B(I)). By comparison the AbdA N-ter, although half the size of that of Ubx, harbors a higher number (4) of smaller regions favoring intra-molecular contacts (Fig. 3B(II)). C-ter sequences display also very different patterns but again the reduced size of the Ubx C-ter limits pattern comparison. This analysis highlights differences in Ubx and AbdA, with a different organization of the N-ter and C-ter sequences and a higher level of compaction of AbdA which may confer a general lower accessibility of AbdA protein sequence for interaction with other proteins. To probe this hypothesis, we calculated accessibility scores for the HX and UA motifs, defined by the sum of interactions formed by the residues of the motif with the rest of the protein (Fig. 3F; WT). Results show that both the HX and UA motifs are less accessible in AbdA than Ubx. Lower accessibility of the HX and UA motifs for interactions with other proteins in AbdA when compared to Ubx may thus contribute to the propensity of motif dispensability for AbdA in vivo functions. Simulation conducted with Ubx and AbdA proteins bearing HX and UA mutated motifs or with Ubx bearing a UA motif of AbdA identity (Ubx^UA(AbdA)^), results in an increase of motifs accessibility, indicating that accessibility is influenced by the HX and UA sequence identities (Fig. 3F).

**Figure 3.**
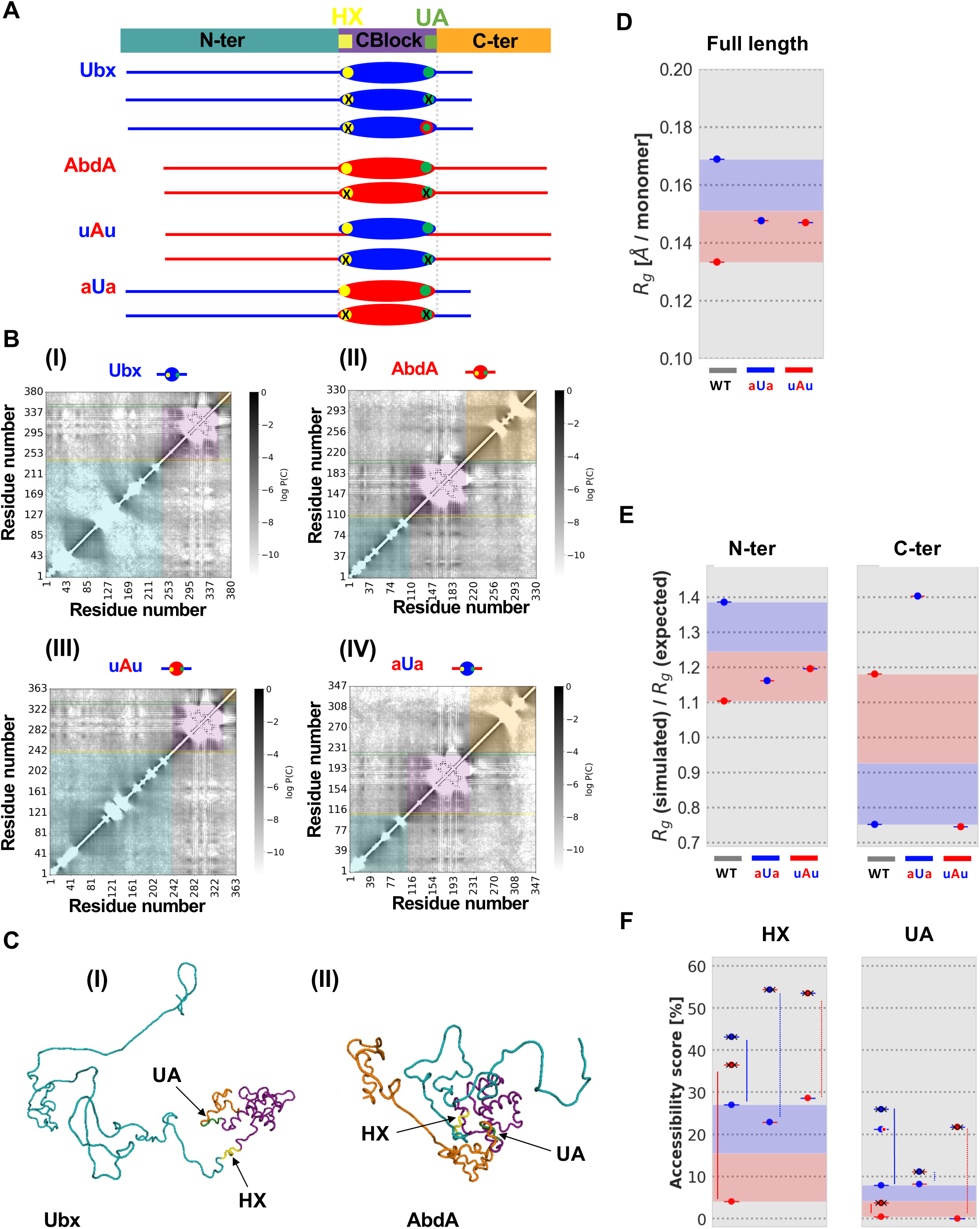
Conformational analysis of Ubx and AbdA from coarse-grained molecular dynamics simulations. **(A)** Schematics of protein regions and chimeras used in the study. The color code representing protein motifs (HX/yellow, UA/green) and protein regions (N-ter/turquoise, CBlock/purple, C-ter/orange) is used in the contact maps and the selected conformation displayed. **(B)** Contact maps of Ubx protein (I), AbdA protein (II), uAu (III) and aUa (IV) proteins. The color scale used in the maps represents the logarithm of the probability of contact formation between pairs of residues, with darker regions indicating a higher contact probability. **(C)** Selected conformations for Ubx (I) and AbdA (II) illustrating the different compactions of Ubx and AbdA. **(D)** Gyration radius of wild type and chimeric Ubx and AbdA proteins. Normalization was according to the size of the protein (Rg expressed in A°/monomer). **(E)** Gyration radius of N-ter and C-ter sequences of wild type and chimeric Ubx and AbdA proteins. Results are presented by the ratio Rg simulated/Rg expected where the Rg expected is calculated according to Flory theory for excluded volume polymers. Ratio of Rg simulated/Rg expected smaller than 1 indicates compact conformations and a ratio above 1 indicates expansion). **(F)** Accessibility (for extramolecular interactions) scores of the HX and UA protein motifs in Ubx and AbdA were quantified based on the sum of interactions formed by the residues of the motif with the rest of the protein. Links between accessibility of wild type and HX and UA motifs mutated proteins (identified by crosses on the protein symbol within the graphs) are highlighted by vertical lines. The change of identity of the UA domain in Ubx (Ubx UA(AbdA)) is represented by a white circled red ball at the UA location.

To further characterize the intramolecular interaction revealed by the Ubx and AbdA contact maps, the associated interaction energies were determined (Fig. 4). We first focused on interaction between protein domains, defining the interaction energies between the central block that includes the HX and UA motifs, and N-ter and C-ter sequences. In both cases, Ubx displays weaker interactions than those engaged by AbdA (Fig. 4A), which may play a significant role in shaping AbdA as a more compact protein, and render the central block that includes the HX and UA motifs less accessible. We next zoomed on the interaction energy between the HX (Fig.4B) or UA (Fig. 4C) motifs, and N-ter, central block and C-ter sequences. For both motifs (HX and UA) and proteins (Ubx and AbdA), the strongest interactions are within the central block. The HX and UA motifs also engage exclusive interactions with N-ter and C-ter sequences respectively, suggesting preferential interaction within a limited distance. Comparing the interaction energy engaged by Ubx or AbdA for each of these interaction classes shows weak differences. Some of these differences may favor a more compact conformation for AbdA (asterisks on Fig. 4B, C).

**Figure 4.**
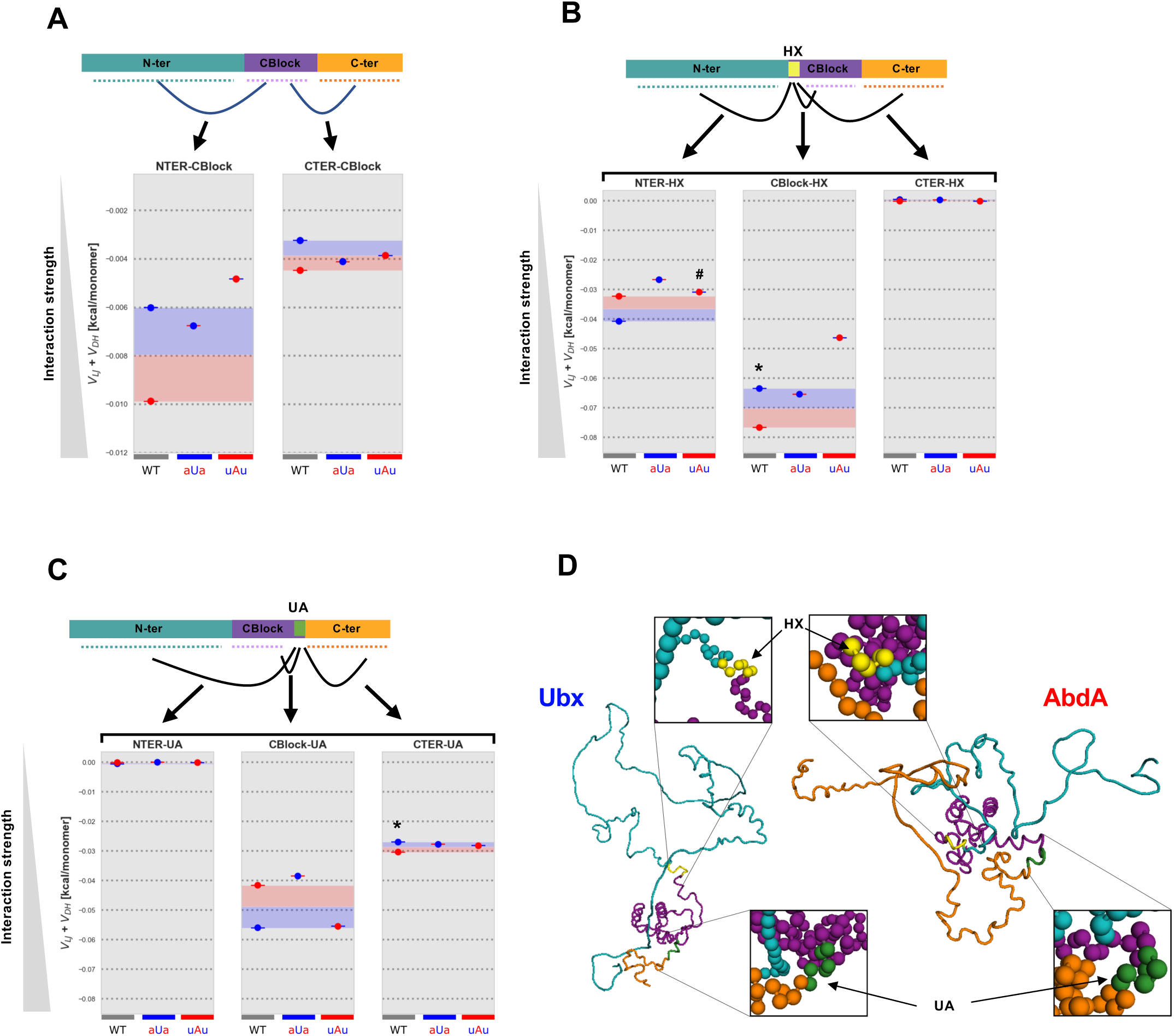
Energetic analysis of the intramolecular interactions in Ubx and AbdA from coarse-grained molecular dynamics simulations. **(A)** Interaction energy between the CBlock and N-ter (left) or C-ter (right) for Ubx, AbdA and the chimeras uAu and aUa. **(B, C)** Interaction energy between the HX (B) or UA (C) motifs with the CBlock, N-ter or C-ter regions. **(D)** Accessibility of the HX and UA motifs illustrated in conformations of Ubx (left) and AbdA (right). The HX motif appears to be the least accessible of the two motifs, possibly because its interaction with the central block is the strongest. This interaction is impacted by the N-ter, as indicated by a similar pattern observed in the N-ter-CBlock interaction. Beads represent amino acids. * highlights cases favoring the view that AbdA is more compact than Ubx. # highlights the only case where the identity of N and C-ter sequences does not guide the strength of interaction.

In summary, coarse-grained molecular dynamics simulations of Ubx and AbdA highlight a dense network of intramolecular interactions, including the HX and UA motifs, illustrated for selected conformation in Fig. 4D. It also suggests that the HX and UA motifs in AbdA are in a protein environment less prone to engage interaction with other proteins, that is globally more compact than Ubx, and in which the central block that includes these motifs engage stronger intramolecular interactions when compared to Ubx.

### Molecular modeling predicts that N-ter and C-ter sequences influence intra-molecular interactions

Coarse-grained molecular dynamics simulations of wild type Ubx and AbdA highlight protein-specific patterns of intra-molecular interactions (Fig. 3B(I) and 3B(II)). These patterns involve interactions within the N-ter, C-ter and central block, but also interactions between these three domains. To assess if/how the N- and C-ter sequences influence Ubx and AbdA intra-molecular interactions, we explored the conformational ensembles of Ubx/AbdA chimeric proteins, where the central block of one protein was flanked by N and C-ter sequences of the other protein (uAu for AbdA central block flanked by Ubx N-and C-ter sequences, and aUa for Ubx central block flanked by AbdA sequences (Fig. 3A). Molecular simulations and subsequent analyses were performed according to the scheme described previously.

Comparison of the contact maps indicates a strong and general interdomain influence. Switching the central block of Ubx with that of AbdA (uAu) strongly affects the pattern of intra-molecular interaction within N-ter sequences, in particular within sequences closer to the central block (compare Fig. 3B(I) and 3B(III)). Changes are also seen in the mirror scenario, where the central block of AbdA was switched with that of Ubx (aUa). In this case, the AbdA N-ter intra-molecular pattern appears simpler (two main domains instead of 4) (compare Fig. 3 B(II) and 3B(IV)). In addition, changes in the pattern within the C-ter sequences are also observed. These changes in intra-molecular contacts modify the gyration radius (Fig. 3D-E). To facilitate the reading of the data, the quantitative graphs were colored in red and blue, with the value range of gyration radius between wild type Ubx and AbdA equally divided in a blue zone, demarcating proximity to Ubx, and a red zone demarcating proximity to AbdA. Remarkably, changes in gyration radius follows the identity of the swapped sequences. Finally, swapping N- and C-Ter sequences influence HX motif accessibility, but not UA motif accessibility, for both Ubx and AbdA (Fig. 3F). As observed in the context of the wild type Ubx and AbdA proteins, mutations of the motifs within chimeras result in an increase of accessibility of the corresponding protein region, again highlighting the contribution of HX and UA sequence identity in defining their level of accessibility (Fig. 3F).

The comparison of the energy of interaction between domains (N-ter/Cblock, C-ter/Cblock) identifies an influence of the identity of the sequences (Ubx or AbdA) on the strength of domain interaction (Fig. 4A). This influence follows the identity of the N and C-ter sequences. Regarding the N and C-ter interaction with the central block, the uAu protein displays a weaker interaction compared to wild type AbdA, which represent a change that mimic Ubx property, where these interactions are weaker compared to AbdA. The mirror modeling using the aUa chimera allows the same conclusions, although the differences in energy observed are within a narrower window (Fig. 4A).

A similar influence of sequences identity was also observed when comparing the energy of interaction between the HX (Fig. 4B) or UA (Fig. 4C) motifs within the central block (HX/Cblock and UA/Cblock) and with the N and C-ter sequences (HX/NTER, HX/C-ter, UA/N-ter, UA/C-ter). Apart from only one exception (see # in Fig. 4B), the identity (Ubx or AbdA) of N and C-ter sequences guides the strength of interaction: for example grafting Ubx N- and C-ter sequences on an AbdA HD results in a protein whose interaction strength are getting closer to that of Ubx; reversely grafting AbdA N- and C-ter sequences on a Ubx HD results in a protein whose interaction strength is closer to that of AbdA. The analysis of the HX or UA interactions within the central block when swapping the N and C-ter sequences further indicates that domain swap also modifies interactions between sequences that have not been changed: this is illustrated by the interaction energies of the aUa and uAu proteins for HX-central block interactions (Fig. 4B, middle panel) as well as of the aUa and uAu proteins for UA-central block interactions (Fig. 4C, middle panel). These observations highlight protein contact interdependence, also apparent in the contact maps (compare turquoise boxes in Fig. 3B(I) and (III) and Fig. 3B(II) and (IV)), including influence from regions that are not modified in sequence.

We conclude that the Ubx and AbdA N and C-ter sequences, strongly divergent between Ubx and AbdA, specify different intra-molecular contacts and contribute to define the overall degree of compactions of the proteins. More specifically, the HX and UA motifs engage interactions with the N and C-ter sequences distinct in Ubx and AbdA. Importantly, modifying N and C-ter sequences not only influences interactions directly involving residues of these protein domains, but also indirectly influences interactions the HX and UA motifs engage within the highly conserved central block.

### In vivo influence of N and C-ter sequence on motif usage

Based on the conclusions reached from molecular modeling, we studied the influence of N and C-ter sequences in routing the use/non-use of the HX and UA motifs, first focusing on AbdA, whose activity in the four in vivo functions studied is largely independent of the HX and UA motifs (Fig. 2). We generated transgenic lines allowing the expression of the uAu chimeric protein (central AbdA block flanked by N and C-ter sequences from Ubx) used in the molecular modeling studies (Fig. 5A). The uAu chimeric protein almost fully loses activity for cerebral branch repression (Fig. S5) and somatic muscle specification (Fig. S6). This generally indicates that the central block does not act as an autonomous functional module and is influenced by surrounding N and C-ter sequences. Importantly uAu retained in vivo activity for beard suppression (Fig. 5B and Fig. S7) and cardiac lineage specification (Fig. 5C and Fig. S8). This allows through mutation of the HX or UA motifs within the chimeric protein environment to question if the Ubx sequences grafted to the AbdA central core influence motif usage. Comparing the impact of HX or UA motif mutations in the uAu protein (beard suppression, Fig. 5B and Fig. S7; cardiac lineage specification, (Fig. 5C and Fig. S8)) to that in the parental AbdA protein, allows addressing if N/C-ter sequences define the Ubx or AbdA type of motif usage (Table S2). Results show that motif contribution is only weakly influenced by Ubx N/C-ter sequences (Fig. 5B’ and Fig. 5C’), with an impact on motif contribution that stays within the AbdA “red” zone for HX, or even for the UA motif diverges from a Ubx type motif contribution. We conclude that for the two available in vivo assays, grafting Ubx N/C-ter sequences to the AbdA central core does not or only weakly allows converting the motif usage mode to that of Ubx.

**Figure 5.**
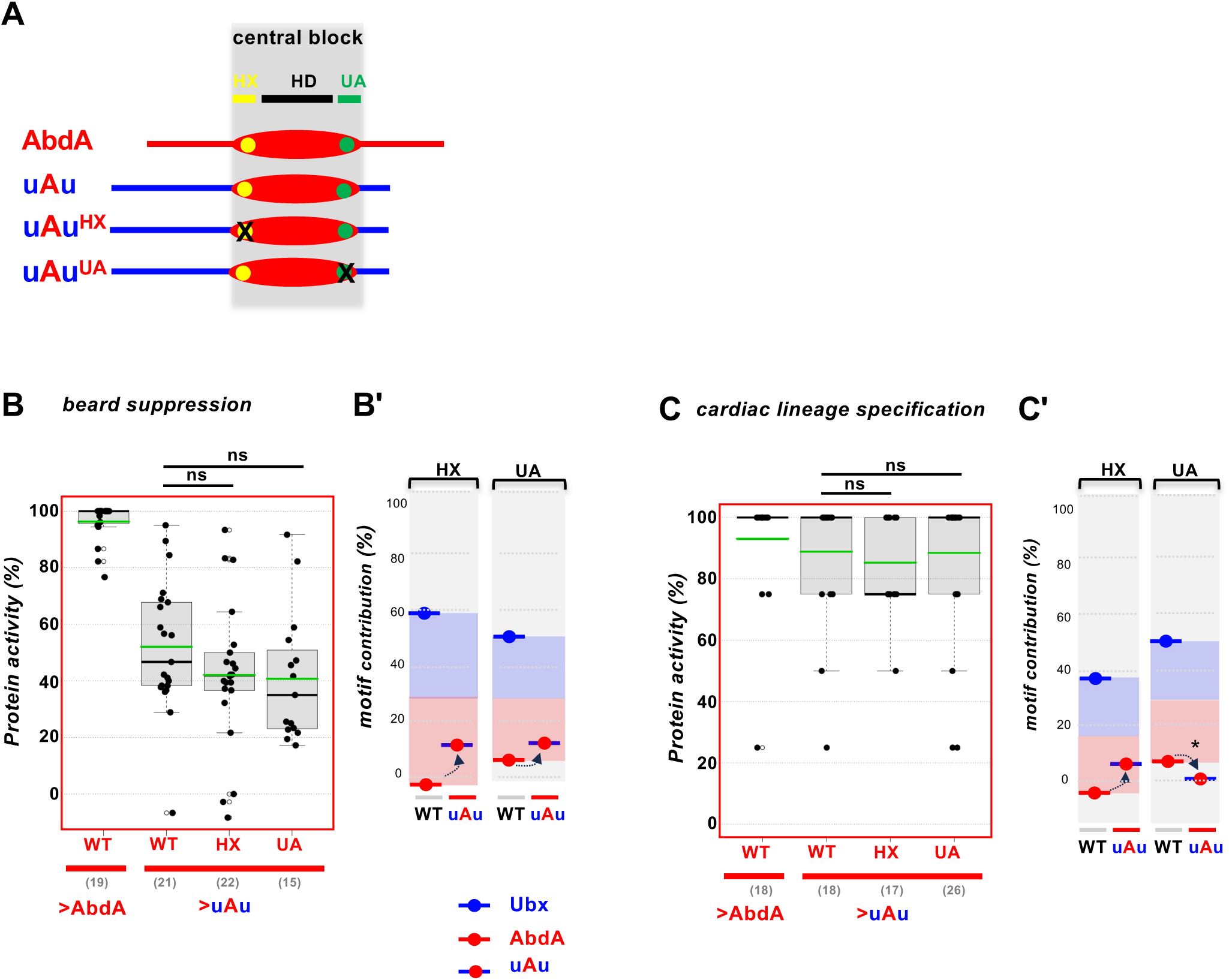
Influence of Ubx N- and C-ter sequences on AbdA motif usage. **(A)** Schematics of chimeric and motif mutated chimeric protein studied. (**B**) Impact of motif mutation on uAu beard repression activity (Fig. S7 and Table S2). **(B’)** Comparison of motif contributions in AbdA and uAu, allowing to visualize the influence of Ubx N- and C-ter sequences on AbdA motif usage for beard repression (arrows). (**C**) Impact of motif mutation on uAu cardiac lineage specification activity (Fig. S8 and Table S2). **(C’)** Comparison of motif contributions in AbdA and uAu, allowing to visualize the influence of Ubx N- and C-ter sequences on AbdA motif usage for cardiac lineage specification (arrows). Grafting Ubx N and C-ter sequences to the AbdA central core generally (with the exception of UA dependency for cardiac lineage specification, see*) weakly affects motif usage by slightly increasing motif dependency for function. Expression (driver used) and readouts are as described in Fig. 2. Mean values are in green. Numbers in brackets on graphs indicate the sample size (N) of each experiment. ns=not significant (p>0,05). Negative values for motif contribution arise when the motif mutated uAu protein activities are higher than motif mutated parental WT proteins.

We applied the same experimental strategy to probe the influence of the AbdA variable N/C-ter sequences on Ubx motif usage, studying the in vivo activity of an aUa chimera protein (central Ubx block flanked by N- and C-ter sequences from AbdA, Fig. 6A). The aUa protein fully loses activity for somatic muscle specification (Fig. S6), impairing further analysis in this specific functional assay, but retained significant activity in the three other functions studied. The activity of aUa^HX^ and aUa^UA^ in cerebral branch suppression (Fig. 6B and Fig. S5) indicates that the mutation of the HX or UA motifs strongly impairs aUa protein activity. On the contrary, motif mutations do not alter aUa protein activity in cardiac lineage specification (Fig. 6C and Fig. S8). The study of beard suppression by the motif-mutated aUa proteins highlights a mixed behavior, with the UA mutation impacting beard suppression while the HX did not (Fig. 6D and Fig. S7). Comparison of the impact of motif mutations within the chimeric and parental proteins allows assessing the influence of N and C-ter sequences (Table S2). Results show that AbdA N/C-ter sequences impose for most functions a switch towards an AbdA type motif usage. The switch is very strong for both motifs in the case of beard suppression (Fig. 6D’) and cardiac lineage specification (Fig. 6C’), where the motif contribution shifts toward the AbdA “red” zone. A moderate impact of N/C-ter sequences was observed for both motifs in the case of cerebral branch suppression (Fig. 6B’). We conclude that grafting AbdA N/C-ter sequences to Ubx central core converts the motif usage mode to that of AbdA.

**Figure 6.**
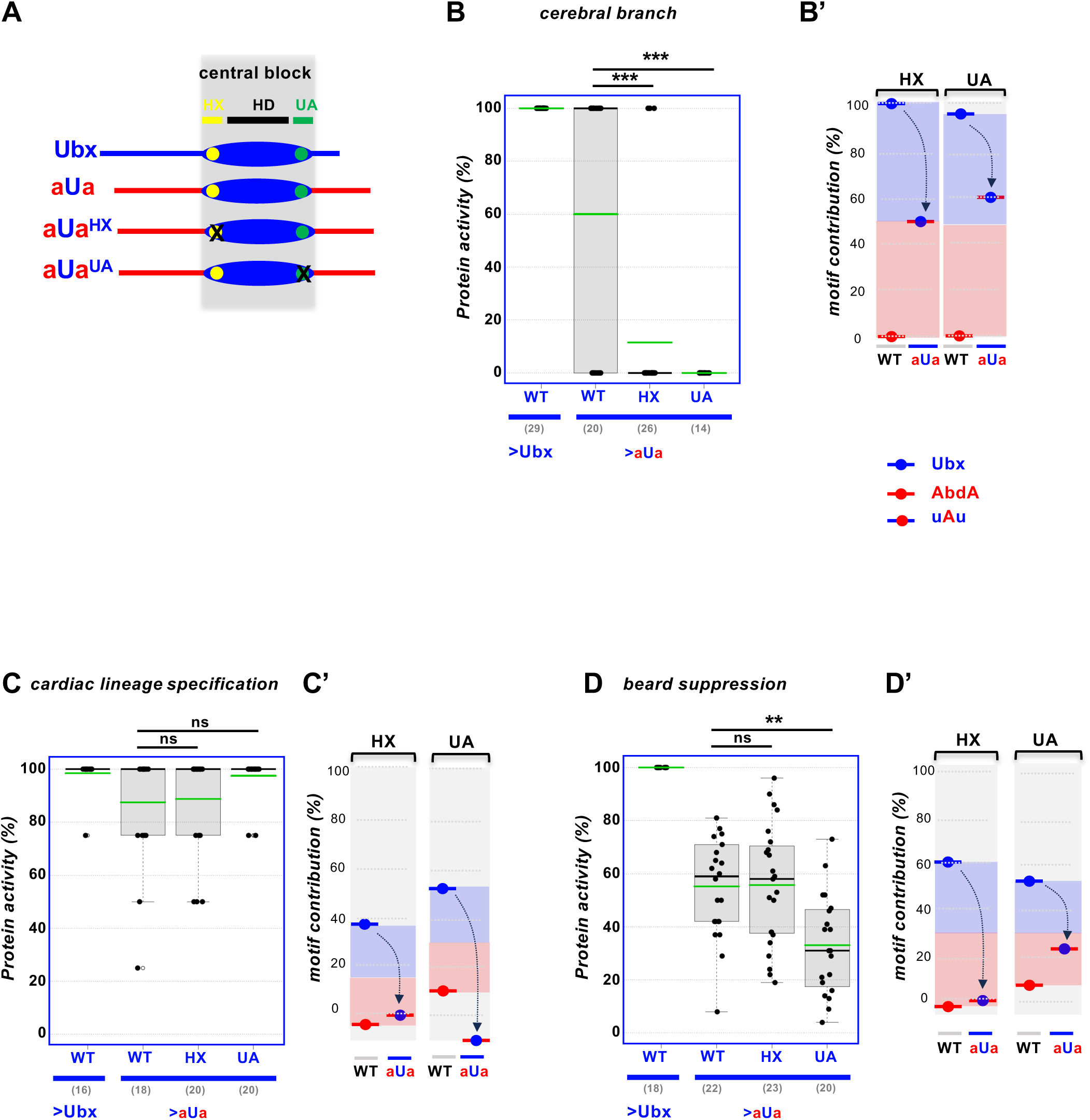
Influence of AbdA N- and C-ter sequences on Ubx motif usage. **(A)** Schematics of chimeric and motif mutated chimeric protein studied. (**B**) Impact of motif mutation on aUa cerebral branch repression activity (Fig. S5 and Table S2). **(B’)** Comparison of motif contributions in Ubx and aUa proteins, allowing to visualize the influence of AbdA N- and C-ter sequences on Ubx motif usage for cerebral repression (arrow). **(C**) Impact of motif mutation on aUa cardiac lineage specification activity (Fig. S8 and Table S2). **(C’)** Comparison of motif contributions in Ubx and aUa proteins, allowing to visualize the influence of AbdA N- and C-ter sequences on Ubx motif usage for cardiac lineage specification (arrows). **(D)** Impact of motif mutation on aUa beard repression activity (Fig. S7 and Table S2). **(D’)** Comparison of motif contributions in Ubx and aUa proteins, allowing to visualize the influence of AbdA N- and C-ter sequences on Ubx motif usage for beard repression (arrows). Grafting AbdA N and C-ter sequences to the Ubx central core confer an AbdA like motif usage type, by decreasing to different extent motif dependency for function. Expression (driver used) and readouts are as described in Fig.2. Numbers in brackets on graphs indicate the sample size (N) of each experiment. Statistical significance using Fisher or Mann-Withney tests (see methods) are *** for 0.0001<p<0.001 and ** for 0.001<p<0.01. ns=not significant (p>0,05). Mean values are in green. Negative values for motif contribution arise when the motif mutated aUa protein activities are higher than motif mutated parental WT proteins.

### Motif usage and N/C-ter sequence influence on Exd-dependent *Dll* repression

We next studied the impact of HX and UA motif mutations on the repression of the limb promoting gene *Distalless* (*Dll)*, an Exd-dependent function shared by Ubx and AbdA and known to rely on the assembly of multi-protein regulatory complexes, involving additional transcription factors, Homothorax (Hth), Sloppy paired (Slp) and Engrailed (En) (Gebelein et al., 2004).

Limbs only form in the thorax and not the abdomen in *Drosophila* and insects as a result of the repression of the limb promoting gene *Dll* in the A1-A7abdominal segments by Ubx and AbdA. The regulation of *Dll* by Hox proteins occurs through direct binding of Hox cofactor complexes on a *Dll* cis regulatory sequences (DME) (Gebelein et al., 2002; Gebelein et al., 2004). Thoracic expression, through ubiquitous *arm-Gal4* driven expression, of either Ubx or AbdA represses *Dll* expression and the activity of the DME-lacZ reporter ((Gebelein et al., 2002), Fig. 7A for Ubx and Fig. S9 for AbdA)). Mean values of the area of fluorescent staining for lacZ-positive cells in the three thoracic segments were scored. Levels of repressive activity were defined by comparing values obtained from WT and Hox variants. Mutation of the HX and UA motifs moderately affects *Dll* repression by Ubx or AbdA: difference in repressive activity was only significant for HX mutation in the case of Ubx, lowering repressive activity from 78 to 66%. In the case of AbdA, significant impacts were seen for both motifs, lowering repressive activity from 95% to 83% (HX mutation) and 79% (UA mutation) (Fig. 7A’ and Fig. S8). The data indicates low and equivalent levels of motifs contributions for the HX motif for both AbdA and Ubx (12%), and a distinct motif contribution for the UA motif, with absence of contribution in the case of Ubx and moderate level of contribution in the case of AbdA (16%), Fig. 7A’’). Strikingly, while in all Exd-independent functions studied, HX and UA were required for Ubx but not (or only very weakly) for AbdA activity, in the case of *Dll* repression, we see either equivalent (HX motif) or stronger contribution, (UA motif) for AbdA activity.

**Figure 7.**
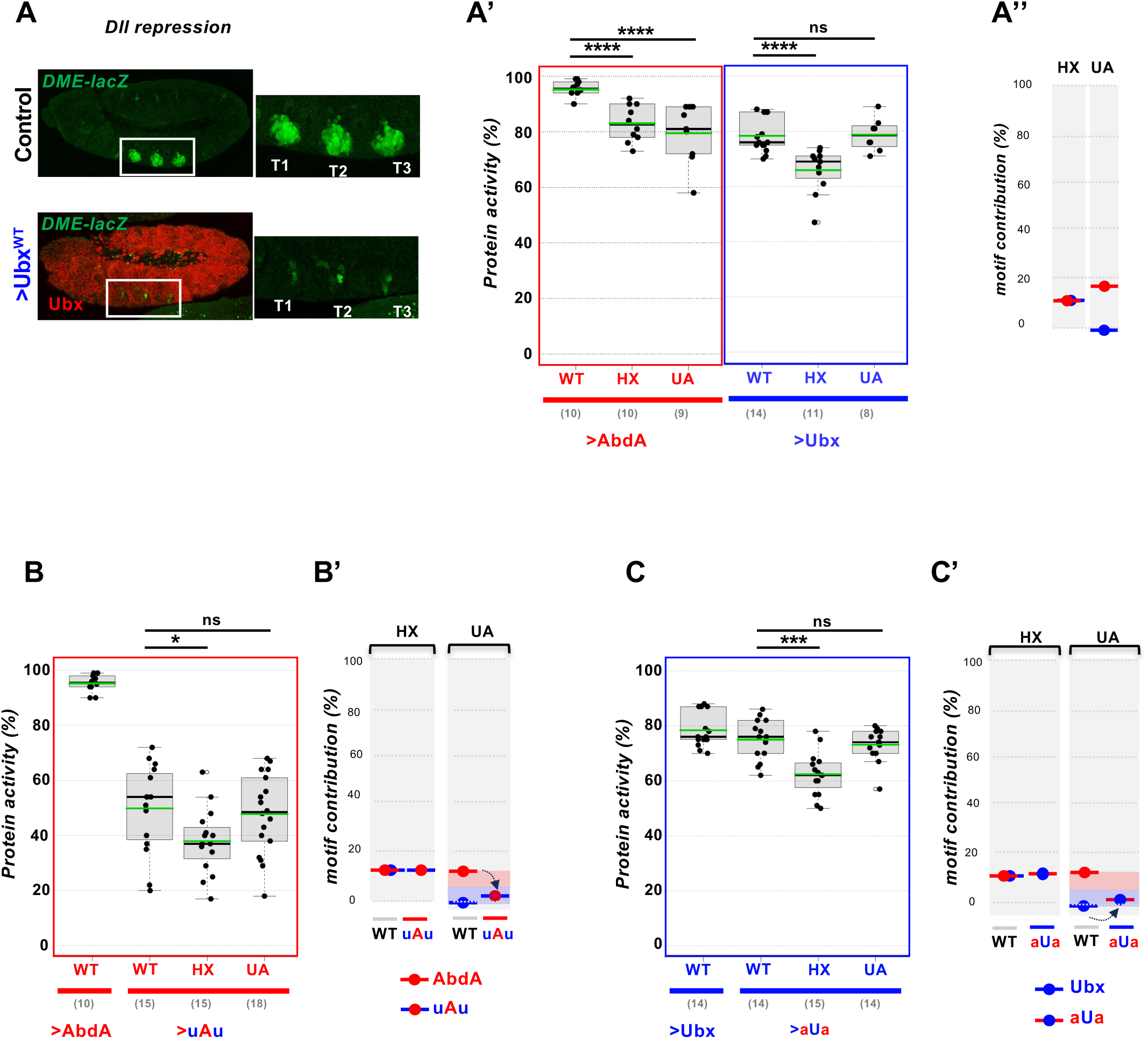
Motif usage for repression of the limb promoting gene *Dll*. **(A)** *Dll* gene expression is confined to the three thoracic segments (T1-3). The DME-lacZ line recapitulates *Dll* expression and regulation (upper panels). *arm-Gal4* driven ectopic expression of Ubx (or AbdA, Fig. S9) strongly represses DME-lacZ expression (lower panel). The area of fluorescent staining for lacZ-positive cells was scored to define protein activities. **(A’)** Protein activities. Repression is less efficient when the HX or UA motifs are mutated in AbdA. In Ubx, only mutation of the HX motif moderately affect *Dll* repressive activity (Fig. S9 and Table S2). **(A’)** Motif contribution was inferred from protein activity (HoxWT activity - HoxMut activity, Table S2). **(B**) Impact of motif mutation on uAu *Dll* repression activity (Fig. S10 and Table S2). **(B’)** Comparison of motif contributions in AbdA and uAu proteins, allowing to visualize the influence of Ubx N- and C-ter sequences on motif usage by AbdA for *Dll* repression (arrow). Grafting Ubx N and C-ter sequences to the AbdA central core alters UA but not HX motif usage. (**C**) Impact of motif mutation on aUa *Dll* repression activity (Fig. S10 and Table S2). **(C’)** Comparison of motif contributions in Ubx and aUa proteins, allowing to visualize the influence of AbdA N- and C-ter sequences on Ubx motif usage for *Dll* repression (arrow). Grafting AbdA N and C-ter sequences to the Ubx central core does not (HX) or only weakly affect motif usage. Numbers in brackets on graphs indicate the sample size (N) of each experiment. Statistical significance using the Mann-Withney test (see methods) are **** for p<0.0001, *** for 0.0001<p<0.001 and * for 0.01<p<0.05. ns=not significant (p>0,05). Mean values are in green.

We next studied the impact of N- and C-ter sequences swap on motif usage. Both uAu and aUa chimeras retained significant *Dll* repressive activity (Fig. 7B, C and Fig. S10), allowing for addressing if N- and C-ter sequences influence UA motif usage. The UA mutated uAu and aUa proteins both exhibit levels of *Dll* repressive activity very similar to that of the parental chimeric proteins (Fig. 7B, C). In the case of Ubx, this is similar to the impact of UA mutation in the wild type protein, resulting in an almost complete absence of motif contribution (Fig. 7C’), and indicating that UA motif usage is not influenced by N- and C-ter sequences swap. In the case of AbdA, the situation is different, as UA mutation impact *Dll* repressive activity: grafting Ubx N- and C-ter sequences to AbdA lowers UA motif contribution to *Dll* repressive activity from 16 to 2% (Fig. 7B’). We thus conclude that UA motif usage in AbdA is influenced by N- and C-ter sequences. For the HX motif, consistent with the lack of differential motif usage, we did not observe any influence of N- and C-ter sequences (Fig. 7B’, C’).

## Discussion

### Distinct motif usage: implication and influence of N- and C-ter IDRs

The present study focuses on how functions are performed by the closely related Ubx and AbdA Hox proteins that share multiple functions (Sambrani et al., 2013; Sambrani et al., 2014). The expectation was that conserved protein motifs, HX and UA, would serve common functions. For four functions examined, beard repression, cerebral branch repression, specification of somatic muscle and cardiac lineage, we consistently observed that while AbdA activity is largely independent of the integrity of the HX and UA motifs, that of Ubx heavily relies on it. Thus, while sharing protein motifs, Ubx and AbdA make distinct use of them to achieve common functions. In one case, the repression of *Dll*, the general requirement of motifs for Ubx and dispensability for AbdA does not apply, highlighting context dependent strategies. The discontinuity between sequence and function conservation observed in most cases suggests that not only motif presence but also motif usage is central to shape protein activity. This calls for caution when using collective insights from protein sequence studies for inferring function for experimentally uninvestigated proteins.

As the main differences in Ubx and AbdA protein sequences lie within IDRs, it suggested that IDRs influence motif usage. The in vivo study of Ubx and AbdA protein swaps where N-and C-ter sequences were permuted was used to assess this influence. In a few situations (3 out of 10, see Fig. 8), swapped proteins were inactive, delineating functional interdependence between the Hox central core and flanking IDRs. This interdependence highlights that a proper match between IDRs and the mostly ordered central core are essential for protein activity, indicative that function is collectively defined by core and IDRs sequences. In the other cases IDRs influence motif usage in a way that matches the origin of the grafted IDR, with influences varying in strength, from strong (dark green in Fig. 8) to weak switches (light green in Fig. 8). In general grafting AbdA N and C-ter sequences to the Ubx central core sequences alleviate the Ubx type motif usage; the reciprocal experiments, grafting Ubx N and C-ter sequences to the AbdA central core sequences by comparison most often only marginally modifies the AbdA-type motif usage, excepting for UA in *Dll* repression. The influence thus appears asymmetric, easily endowing Ubx with AbdA properties, but hardly conversely. This asymmetry may reflect that swapping domains easily disrupts interactions and function, but more hardly create novel functionally relevant interactions.

**Figure 8.**
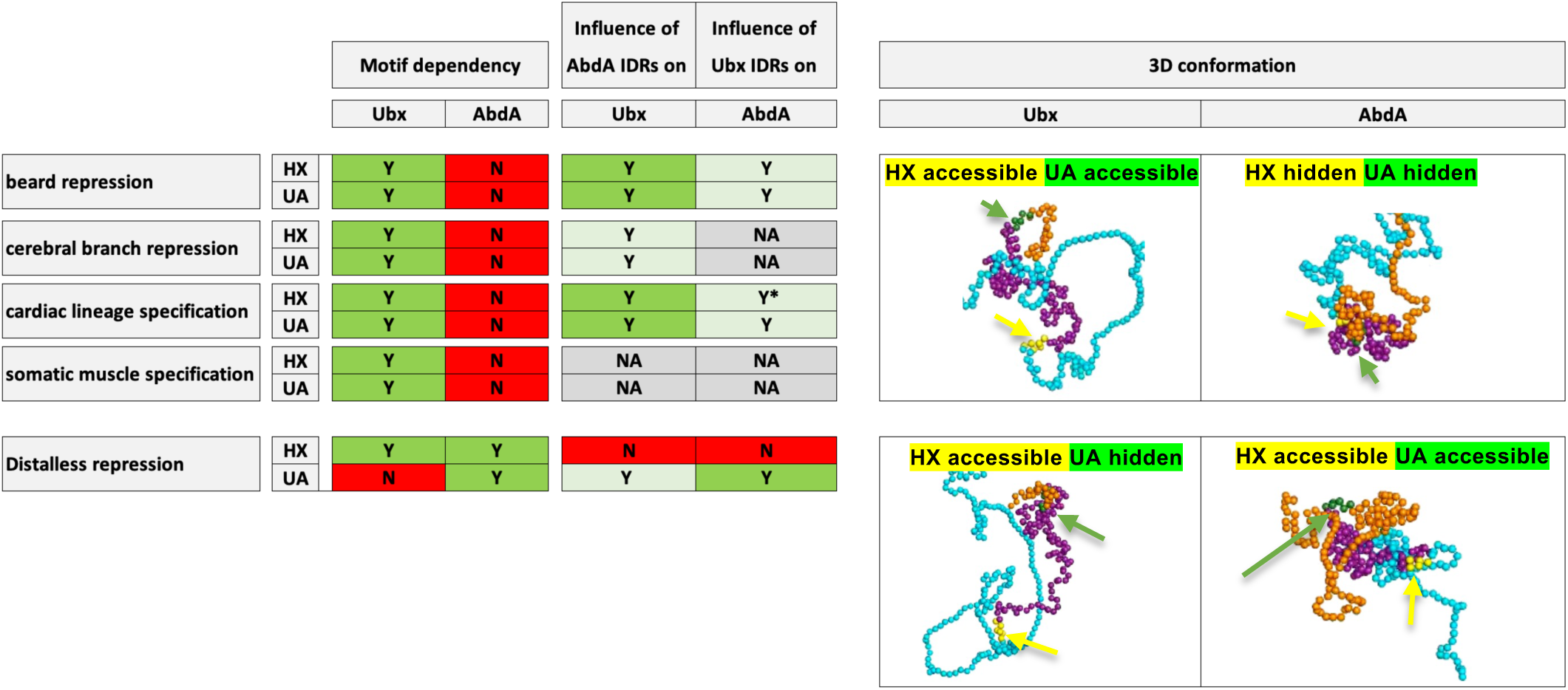
Motif dependency, influence of IDRs and suggested links to protein conformation. **Motif dependency (left panels).** Motif dependency on HX and UA motif usage for each of the five in vivo functions studied are summarized. Y (=Yes) within green boxes indicate motif dependency, N (=No) within red boxes indicate absence of motif dependency. With the exception of *Dll* repression, Ubx consistently display HX and UA motif dependency while AbdA does not. **Influence of AbdA IDRs on Ubx motif usage and of Ubx IDRs on AbdA (middle panels).** Y (=Yes) within green boxes indicates influence and N (=No) within red boxes the absence of influence. NA= Not Assessed as the uAu or aUa proteins are inactive in the biological readout considered. Dark green boxes highlight strong switches, light green boxes weak switches. Switches were qualified as strong when motif contribution of the chimera gets closer to that of the protein that provides N-and C-ter sequence, highlighted in Fig. 5-7 by a change from the red to blue (or reverse) zone. Weak switches are changes that remains closer to the protein that provides the central core, highlighted in Fig. 5-7 by a change from the red to blue (or reverse) zone. With the exception of *Dll* repression, AbdA IDRs generally have a stronger influence on Ubx activity than Ubx IDRs on AbdA activity. * highlights the single case where activity change is not towards the identity of the graft (see Fig. 5C’). **Snapshots of the molecular modeling simulations consistent with motif dependency except for *Dll* repression (upper right panels).** A Ubx conformations with an HX and UA accessible motifs and an AbdA conformation with hidden HX and UA motifs would account for the motif dependency observed for beard repression, cerebral branch repression, cardiac cell lineage specification and somatic muscle specification. These conformations are frequent within the conformation ensemble revealed by molecular modeling simulations. **Snapshots of the molecular modeling simulations consistent with motif dependency for *Dll* repression (lower right panels).** A Ubx conformation with an HX accessible and UA hidden motifs and an AbdA conformations with HX and UA motifs accessible would account for the motif dependency observed for *Dll* repression. These conformations are infrequent within the molecular modeling simulations. To occur they likely rely on protein interaction within a larger regulatory complex.

### Bases for distinct motif usage and its control: hypotheses from in silico molecular modeling

To start identifying bases for differential motif usage, we used in silico molecular modeling of Ubx and AbdA proteins to identify intrinsic properties that may contribute to distinguish the activity of the two proteins. The analyses identify large and dynamic sets of intramolecular interactions which certainly largely shape the 3D folding of Ubx and AbdA. One consequence of these intramolecular interactions is that AbdA seems to fold in a more compact protein than Ubx, resulting in a reduced accessibility of the HX and UA motifs for interaction with other proteins. In four of the five studied in vivo functions we found a correlation between motif accessibility defined *in silico* and motif usage deduced from in vivo functional analyses (Fig. 8): the dispensability of the HX and UA motifs for AbdA in vivo function aligns well with the compact nature of AbdA for which the HX and UA motifs accessibility is low; conversely the higher accessibility of the motifs in Ubx is more prone to interaction with other proteins, possibly explaining requirement for Ubx function.

The fifth in vivo functions studied, the repression of *Dll*, stands out since it does not follow the general requirements trend. Although, the *Dll* case appears unique in our study, it may illustrate a large cohort of Ubx and AbdA downstream targets. Interestingly, *Dll* repression has been extensively studied (Gebelein et al., 2004; Merabet et al., 2007; Noro et al., 2011; Saadaoui et al., 2011), and shown to rely on the assembly of quadripartite regulatory complexes on *Dll* cis regulatory sequence: Ubx/Exd/Hth/En and AbdA/Exd/Hth/Slp. We propose that within such large regulatory complexes, protein-protein interactions impose constraints and conformations that cannot be predicted by molecular modeling of isolated proteins. These constraints may favor conformations that in the monomeric modeling are rather rare or even inexistant, as suggested by IDRs acquiring specific folding in contact with other protein, usually in contact with structured protein domains (Banerjee et al., 2024; Holehouse and Kragelund, 2024; Tompa et al., 2014; Wang and Zhang, 2025).

We also used in silico molecular modeling to ask if intrinsic Ubx and AbdA properties help understanding how IDRs influence motif usage. In the case of the HX motif, the study of domain swapped proteins in silico supports the view that, as observed in vivo, the identity of IDRs influences HX motif accessibility: grafting AbdA IDRs to the Ubx central block reduces predicted accessibility while the symmetrical graft (Ubx IDRs to the AbdA central block) increases it. We however note that the strength of influence observed in silico, stronger for Ubx IDRs when compared to AbdA IDRs, does not correlate with the strong influence of AbdA IDRs and weak influence of Ubx IDRs observed in vivo. In the case of the UA motif, no influence of IDRs was observed in silico, perhaps as a result of weaker predicted accessibility when compared to the HX and of its propensity to fold as a continuation of the HD recognition helix (Foos, 2015; Ortiz-Lombardia et al., 2017). This forbids any parallel with the in vivo observed influence of IDRs on UA motif usage.

We concluded that in silico molecular modeling on isolated Ubx and AbdA proteins that grasps intrinsic protein properties can only predict some aspects of molecular mechanisms at play. Higher order molecular modeling, including DNA and additional protein partners, and the identification of functionally relevant compositional differences between Ubx and AbdA IDRs, should provide a deeper understanding of how IDRs control protein activity.

## Material and Methods

### Fly stocks and transgenic lines

The following fly stock were used: w^1118^ (as control); *24B-Gal4* (BL 1767); *btl-GAL4; arm-Gal4; svp-lacZ* (BL 26669); *DME-lacZ* (Gebelein et al., 2002), UAS-Ubx (Merabet et al., 2007); UAS-Ubx^HX^ and UAS-Ubx^UA^ and UAS-Ubx^UA(AbdA)^ (this study); UAS-abdA, UAS-abdA^HX^ and UAS-abdA^UA^ (Merabet et al., 2011). The UA centered following changes were introduced in Ubx for generating the Ubx^UA(AbdA)^: <u>IQ</u>**A<u>I</u>KE<u>L</u>NEQ**<u>EK</u> to <u>LR</u>**A<u>V</u>KE<u>I</u>NEQ**<u>AR</u> (UA motif in bold). Ubx and AbdA modified cDNAs were cloned into pUAST and injected by Bestgene in w^1118^ strain. HX and UA variants are defined in Fig. 1A. aUa and uAu Chimeras were produced by PCR from WT *Ubx Ia* and *abdA* cDNAs. aUa and uAu chimeras contain a HD central block from Ubx or AbdA respectively, starting with HX motif and ending with UA motif (see Fig. 1A), flanked by N and C-terminal sequences from AbdA for aUa and from Ubx for uAu. For chimeras bearing motif mutations, the central block was amplified from cDNAs containing the suited Ubx or AbdA mutations. Constructs were HA tagged at their C terminus and cloned into pUAST attB using the In-Fusion HD cloning kit (Takara Bio USA). PhiC31 injections were done by Bestgene in the Bloomington 9752 strain. Primer sequences used are described in Table S3.

*Ubx* and *AbdA* HA tagged loci were generated by CRISPR mediated Homologous Directed Recombination (HDR). In both cases, the last exon was targeted by HDR to introduce a 3XHA tag and a selection cassette. The selection cassette consists of a 3xP3-dsRed eye reporter gene, flanked by two loxP sites. The donor plasmid sequences used for HDR were designed in silico and synthesized by GenScript (Fig. S11). gRNAs were cloned into the pCFD4-U61-U63 tandem gRNA plasmid following the CRISPRflydesign protocol (http://crisprflydesign.org/wp-content/uploads/2023/12/Cloning-with-pCFD4.pdf ; see Table S3 for oligo sequences). Both plasmids were injected into approximately 500 nos-Cas9 (BDSC 54591) preblastoderm embryos (100 ng/ml of sgRNA plasmid + 500 ng/ml of donor plasmid). G0 males were then individually crossed to third chromosome balancer females. G1 male progeny with red fluorescent eyes were selected and individually crossed to third chromosome balancer females to establish an isogenized stock with the modified chromosome.

### Crosses, egg collections, cuticle preparations, immunostaining and in situ hybridization

Ubx and AbdA wild type and variant protein expression were obtained by crosses with Gal4 drivers. Cross temperatures were adjusted to reach expression levels similar to endogenous Ubx and AbdA proteins (+/-15%; see (Merabet et al., 2011; Merabet et al., 2007) and below in the following section for a detailed description of the procedure). Egg collections, cuticle preparations, immunostaining and in situ Hybridization were performed according to standard procedures. The *nau* digoxigenin RNA-labeled probe was generated by in vitro transcription from a cloned cDNA. Primary antibodies used were mouse anti-Ubx (1/100, FP3.38); rabbit anti-AbdA P48 (1/200, Yacine Graba); rabbit anti-Dmef2 (1/2000, gift from L. Perrin); rat anti HA (1/500, Roche); rabbit anti-β-galactosidase (1/500, MP Biomedical); chicken anti-β - galactosidase (1/1000, Abcam); chicken anti-GFP (1/1000, Aves); anti-digoxigenin coupled to biotin (1/500, Jackson ImmunoResearch). Secondary antibodies used were coupled to AlexaFluor 488, AlexaFluor 568 (1/500, Life Technologies), or to biotin (1/500 Jackson ImmunoResearch).

For the illustration of Fig. 1D, embryos were stained for Ubx and AbdA. As previously described, Ubx is predominantly expressed in the anterior and AbdA in the posterior compartment (Gebelein et al., 2004). To highlight the overlapping expression patterns, images were treated as follows: projection of confocal Z sections was processed in Photoshop by applying a negative filter to each staining. Blue and red colors were assigned to Ubx and AbdA respectively. The embryo used for Fig. 1D is displayed in Fig. S12.

#### Quantification of UAS-driven Ubx and AbdA proteins (WT, HX, UA and UA(AbdA))

Gal4 individuals were crossed with heterozygote/hemizygote UAS-Hox lines at different temperatures (18°C,22°C,25°C and 29°C). Given the hetero/hemizygotie of the UAS-Hox lines, part of the progeny activated the UAS-Hox transgene, while the other part was not. Embryo were collected overnight (0-12h collection), fixed and immune-stained according to standard procedures. Anti-Ubx (mouse Mab FP3.38, DSHB AB_10805300) and anti AbdA (rabbit polyclonal P48, Yacine Graba) were used.

To select temperature conditions where UAS-driven Hox proteins are expressed close to endogenous protein levels (+/-15%), embryos expressing the UAS-Hox transgene (thoracic and abdominal expression) or not (abdominal restricted expression) were identified on the same slide. Levels of endogenous Hox expression was deduced from quantification of Hox staining in embryos with abdominal restricted Hox expression, and levels of UAS-driven expression was deduced from quantification of Hox staining in thoracic regions (where only UAS-driven Hox is present) of embryos with unrestricted Hox expression. An identical area zone was used in the abdomen of embryos with restricted Hox expression (endogenous level), and in the equivalent region in the thorax of embryos with ubiquitous expression (UAS-driven expression) to define the mean intensity of the staining (Fiji/ImageJ>polygon selection tool> analyse > measure > mean intensity). The positioning of this area zone depended on the readout of the experiment (Fig. S13).

Following this procedure, we found that temperature conditions to approximate endogenous expression levels were not dependent on the driver used (ie tissue targeted by the expression) but on the identity of the UAS-Hox line. The following temperatures allowed to stay within the “+/-15 %” range irrespective of the Gal4 driver used: UAS-Ubx^WT^, 22°C; UAS-Ubx^HX^, 29°C; UAS-Ubx^UA^, 29°C; UAS-Ubx^UA(AbdA)^, 29°C; UAS-AbdA^WT^, 25°C; UAS-AbdA^HX^, 25°C; UAS-AbdA^UA^, 25°C.

To further compare endogenous Ubx and AbdA expression levels, we used Ubx and AbdA CRISPR lines engineered fly lines where the endogenous Ubx and AbdA loci were modified so to fuse an HA tag at the C-terminus of each protein (Fig. S11) as a proxy. The generated CRISPR lines accurately reproduce the Ubx and AbdA expression patterns (Fig. S11). Quantification of HA staining in CRISPR lines was performed within A1 for Ubx, and in A2 for AbdA, avoiding the segment borders in both cases. Results indicate that endogenous Ubx and AbdA, assessed using the same HA antibody in the same experimental conditions, are expressed at similar levels (Table S4, Fig. S11). As endogenous Ubx was used to set the level of UAS-driven Ubx proteins, and similarly endogenous AbdA was used to set the level of UAS-driven AbdA proteins, we concluded that UAS-driven Ubx and AbdA proteins are expressed at similar levels.

### Quantification of UAS-driven Ubx and AbdA chimeric proteins (aUa, aUa^HX^, aUa^UA^, uAu, uAu^HX^, uAu^UA^)

Lines to express UAS-driven Ubx/AbdA chimeric proteins were generated using site directed genomic insertion (site VK37 in BDSC stocks 9752). The resulting transgenes were thus all located at the same genomic position (22A3) minimizing possible expression differences owing to the influence of the local genomic environment.

Temperature conditions allowing for uAu chimeras (WT, HX or UA mutated) expression close (+/-10%) to endogenous expression level were selected according to the same principle described for assessing expression levels of UAS-driven Ubx and AbdA proteins. In brief, homozygote arm-Gal4 individuals were crossed with heterozygote UAS-uAu lines at different temperatures (18°C, 22°C, 25°C and 29°C). Embryos were simultaneously stained with the anti-Ubx (mouse Mab FP3.38, DSHB AB_10805300), that recognize an epitope within the Ubx N-ter sequence, thus recognizing all uAu chimeras and with an anti-HA. Confocal images were acquired with identical settings. Intensities from anti-Ubx staining were measured either in A1 for wild-type embryo (endogenous level) or in thoracic segments in embryos ectopically expressing uAu chimeras (transgenic level). uAu transgenes expression at 29 °C were found to be the closest (+/-10%) to endogenous Ubx levels. Intensities values from anti-HA staining were also measured in thoracic segments of embryos ectopically expressing uAu chimeras. We found that these values are approximately the same (+/-10%) at 29 °C for the three uAu chimeras (WT, HX or UA mutated).

The mirror experiments using UAS-aUa (WT, HX or UA mutated) transgenic lines was not possible since the AbdA antibody used does not recognize aUa proteins. In these instances, embryos were only stained the HA antibody. Since we have established that intensity values resulting from anti-HA staining following uAu expression at 29 °C is close to endogenous Ubx, we used these values as reference to adjust UAS-aUa expression. Measurement of anti-HA staining in thoracic segments following aUa expression at 29 °C shows values similar to uAu expression at 29 °C. This is consistent with a unique genomic location of the transgenes.

#### Quantifications of in vivo protein activities

Levels following induced expression were quantified and experimental conditions ensuring close to endogenous levels were selected. Quantification of protein activities are according to (Merabet et al., 2011) that describes the effects of AbdA motif mutations, with minor modifications (see below). For each function examined, at least ten embryos or larvae were used to score the activity of each protein. Boxplot were drawn using the R software. Medians are displayed. Statistical significance was assessed by Fisher tests for data pertaining to cerebral branch repression, and Mann/Withney test for data pertaining to beard suppression, cardiac lineage, somatic muscle specification, and *Dll* repression. Motif contributions for each protein and each in vivo functional readout were defined as the difference of protein activity in the wild type and motif-mutated protein. When the mutated protein is more active than the wild type protein, the motif contribution score is negative. This appears to be the case in a few instances (see Table S2).

##### Cerebral branch (CB) repression

Level of protein repressive activity on CB formation was followed by the expression of UAS-GFP driven by *btl-GAL4* (trachea specific driver). Presence or absence of CB was scored. Absence of CB formation corresponds to 100% of protein activity; 0% of protein activity correspond to presence of the CB. Embryos in lateral views were considered. Scoring always concerned one hemi-embryo.

##### Somatic muscle formation

Embryonic somatic muscle formation is followed by the expression of *nau*. The number of ectopic ventral cluster in TI, T2, T3 and A8 was scored. Full protein activity (100%) is when 4 additional ventral clusters are induced, 75% is for 3 additional clusters, 50% for 2, 25% for 1 and 0% when no ectopic ventral cluster is seen. Embryos in lateral views were considered. Scoring always concerned one hemi-embryo.

###### Cardiac cell lineage specification

The *svp-lacz* reporter gene allows delimiting segments in the heart. The number of cardiac cells in the two posterior most segments of the anterior aorta (in each hemi-embryo), ranging from 8 to 12, was scored. Cardiac cells were counted only in the two posterior most segments of the anterior aorta as the most anterior part of the heart dives inside the embryo in a region that is *svp* positive, making it difficult to unambiguously identify the 6 cell lineage transformation in that region. A full transformation results in 12 cells (instead of 8 in the WT). Proteins inducing this type of transformation were assigned a score of 100% activity. Partial transformations, ranging from 11, 10, 9 to 8 cells were assigned protein activities of 75%, 50%, 25 and 0%.

##### T1 beard suppression

T1 beard denticles were scored in first instar larvae using the ImageJ cell counter. On average, the T1 beard of a wild type embryo is composed of 180 denticles. Protein activity was defined as follows: 100-100x(#denticles of ectopic expressing Hox)/(#denticles in control larvae). 100% of activity is a complete beard repression. Values between 0 and 100% correspond to partial suppression of the beard.

###### Distalless DME repression

Levels of *Dll* repressive activity was followed by the expression of the *DME-lacZ* reporter (Gebelein et al., 2002; Gebelein et al., 2004). Mean values of the area of fluorescent staining for lacZ-positive cells in the three thoracic segments using the ImageJ polygon selection tool were scored. Protein activity was defined as follows: 100-100x(mean value of ectopic expressing Hox embryo)/(mean value in control embryo). Embryos in lateral views were considered. Scoring always concerned one hemi-embryo.

#### Coarse-grained molecular dynamics simulations

To investigate the conformational ensemble of the Hox proteins, we performed coarse-grained molecular dynamics simulations of Ubx, AbdA, chimeric proteins (aUa and uAu), HX and UA mutated chimeras, and Ubx^(UA(AbdA))^. Each residue is represented by a bead centered on the *C_α_* atom. The general potential energy function (**Eq. 1**) comprises several terms: The first three terms describe "bonded" interactions: harmonic bonds, harmonic angular geometry, and backbone rotation around dihedral angles. The remaining terms describe "non-bonded" interactions: native and disordered contacts, repulsions, and electrostatic interactions using the Debye-Hückel potential.

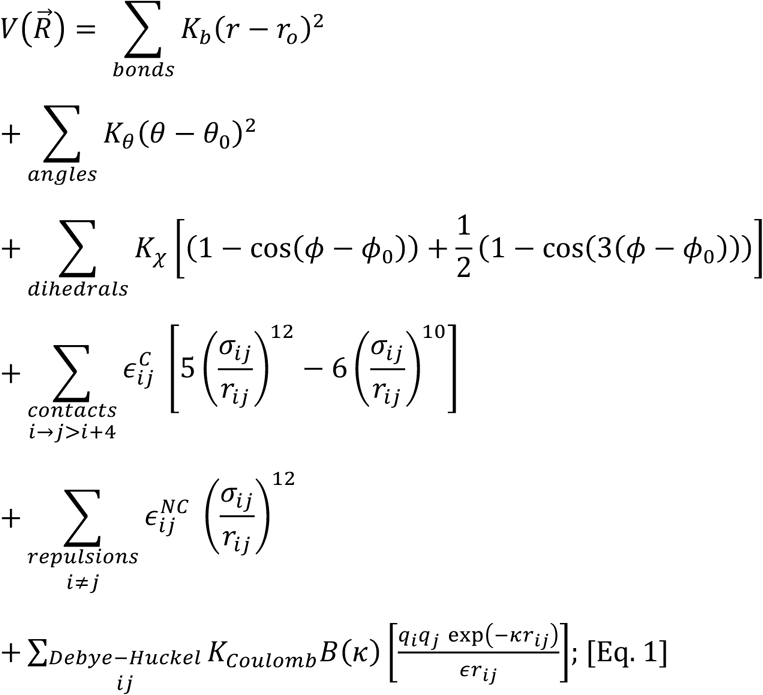

Where *K_b_*, *K_θ_*, *K***_z_** are constants defining the stiffness of the bonded potential. For the ordered regions, the potential constants used were *K_b_* = 100 *kcal mol*^−1^, *K_θ_* = 20 *kcal mol*^−1^, *K***_z_** = 1 *kcal mol*^−1^ . *θ* and *φ* are angles in radians, *r_i_*_j_ represents the interbead distance in Å. *ε_i_*_j_ and *σ_i_*_j_ define the shape of the 12-10 Lennard-jones potential, with the former representing the well depth and the latter the van der Waals radius. In the Debye–Hückel potential, *q_i_* and *q*_j_ are the charges of the residues at the respective indices, ε is the solvent dielectric constant, and *K_Coulomb_* = 4*πε*0 = 332 *kcal*/*mol*, *κ* is the screening factor and was set to 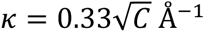, where C is the salt concentration in molar. For dilute solution, the salt-dependent coefficient, is approximately *B*(*κ*) ≈ 1.

Initial structures were generated with AlphaFold using the ColabFold server. Native contacts were identified using the CSU algorithm for the homeodomains of Ubx and Abda, and modeled with a potential constant of 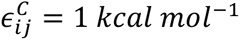. Pairs not defined as native contacts were treated as repulsions 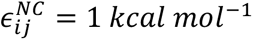. In order to represent the higher conformational flexibility of IDRs, the bond angles stiffness parameter *K_θ_* was set to half the value used for ordered regions (10 *kcal mol*^−1^), dihedrals angles were removed, and contacts were defined for interacting pair (*i*, *j*) within the IDR or between the IDR and the homeodomain where *j* > *i* + 4 according to the FB-HPS model values for *ε_i_*_j_ and *σ_i_*_j_ (Dannenhoffer-Lafage and Best, 2021).

The systems were simulated using Langevin dynamics for a total of 10^7^steps at a temperature of 0.4 (reduced units). The dielectric constant was set to 80 and the salt concentration to 0.02 M. All the analyses are based on three independent simulations for each protein.

#### Probability of contacts

To start deciphering the pattern of interaction between the different regions in each protein we calculated the probability of contacts for each residue pairs. The total number of contacts for each pair was determined in a binary fashion based on a cutoff distance of 1.2 *σ_i_*_j_ . Interacting pair at a distance *r_i_*_j_ below the cutoff were considered “in contact”, while pairs above the cutoff were considered “not in contact”. The probability of contact was obtained by dividing the sum of contacts by the total number of frames.

#### Radius of Gyration

The compaction of the proteins or the selected regions were quantified by the *R*_g_ values, and calculated according to the definition:

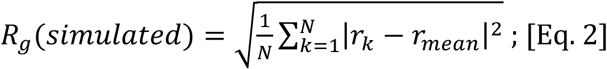

Where N is the length of the protein, *r_k_* is the position of residue k, and *r_mean_* is the mean position of the protein. Theoretical values for each protein were established using Flory’s seminal power law:

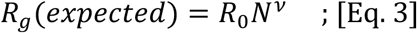

Where *N* is the number of residues, *R*_0_ is a prefactor and *ν* isFlory’s exponent. The value for the constants were *R*_0_ = 0.254 and *ν* = 0.52.

#### Interaction energy

To further characterize the dense network of intramolecular interactions, the energy of interaction was calculated for each frame using the definition of the nonbonded terms (Eq. 1). In each case, only the relevant interaction pairs (*i*, *j*) were taken into account.

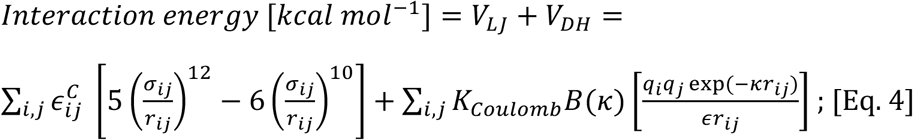

#### Accessibility Score

To correlate molecular simulations with the importance of motifs identified from experimental results, we defined an accessibility score for each motif, *A_i_*. The accessibility score was quantified based on the sum of interactions, formed by the residues of the motif with the rest of the protein. An interaction was defined as any inter-residue distance (i.e., any bead within the motif interacting with any bead outside of it) of less than 10 Å. The resulting sum, *M_i_*, was then averaged across all frames and normalized by the protein length. Finally, the accessibility score for each motif 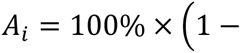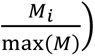, was obtained by scaling the respective *M* between 0 and 1 with respect to the motif with the highest number of interactions.

## Acknowledgement

Work in the YG group is supported by the CNRS and Aix Marseille University. This work received support from the French government under the France 2030 investment plan, as part of the Initiative d’Excellence d’Aix-Marseille Université -A *MIDEX" AMX-22-RE-AB-152., and the FRM (EQU202403018061). MM was supported by a MESR fellowship, LM by an AMIDEX fellowship. Y.L was supported by the Israeli Science Foundation (2072/22) and by a research grant from the Estate of Gerald Alexander. We thank DSHB for antibodies and the Bloomington stock centers for fly lines. All material is available upon request.

