## Supplemental info for "Intrinsically disordered regions influence protein motif usage in *Drosophila* Ubx and AbdA Hox proteins"

### *cerebral branch suppression*

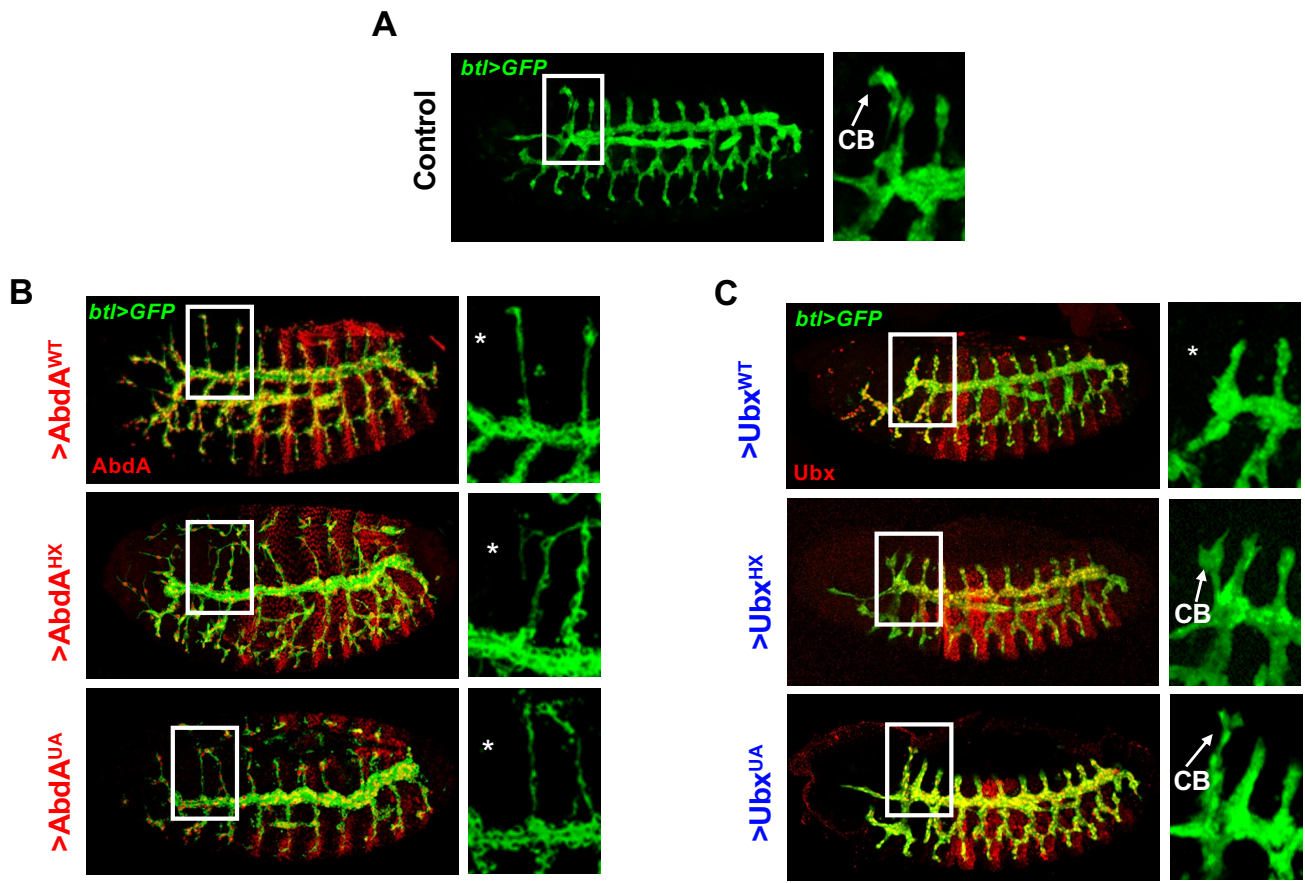

**Figure S1. Ubx and AbdA motif dependency for cerebral branch repression.** Trachea are visualized by *btl-Gal4* driven GFP (green). **(A)** In control embryos cerebral branch (CB) forms exclusively in T2 (arrow). **(B)** Ectopic expression of *AbdA<sup>WT</sup>*, *AbdA<sup>HX</sup>* and *AbdA<sup>UA</sup>* variants (red) suppress CB formation (asterisk). **(C)** Ectopic expression of *Ubx<sup>WT</sup>* suppress the formation of the CB (asterisk) while *Ubx<sup>HX</sup>* and *Ubx<sup>UA</sup>* (arrows) do not. Right panels are enlargement of white boxes.

#### *somatic muscle specification*

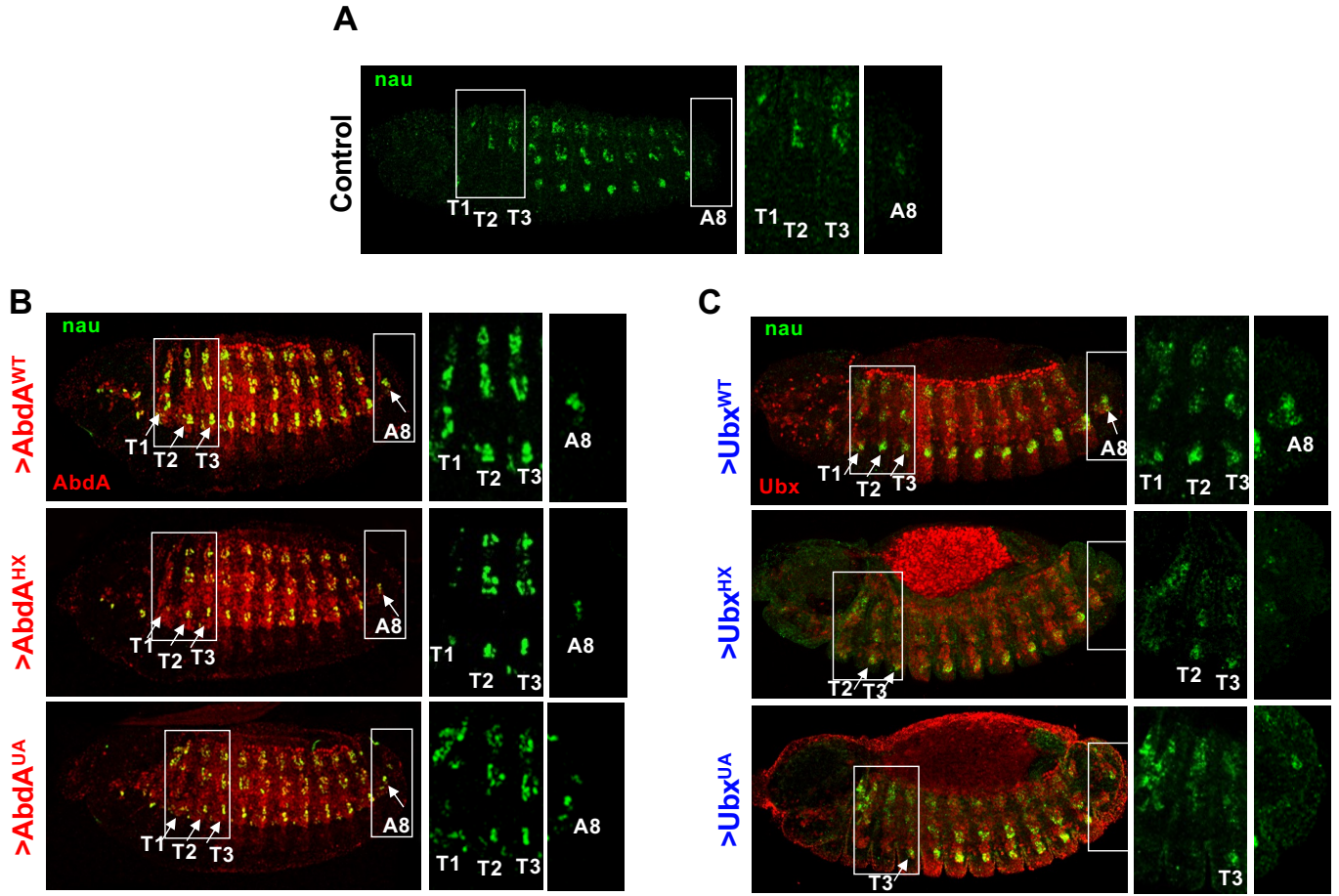

**Figure S2. Ubx and AbdA motif dependency for ventral somatic muscle specification.** Somatic muscle formation is visualized by the expression of *nautilus* (*nau*, green). **(A)** in control embryos thoracic segments T1-T3 and Abdominal segment A8 lack a ventral somatic muscle cluster. **(B)** *24B-Gal4* mesodermal expression of *AbdA*<sup>WT</sup> (red) induces ectopic somatic muscles ventrally in T1-T3 thoracic segments and in A8 (arrows). This ability is intact for *AbdA*<sup>HX</sup> and *AbdA*<sup>UA</sup>. **(C)** *24B-Gal4* mesodermal expression of *Ubx*<sup>WT</sup> (red) induces ectopic somatic muscles ventrally in T1-T3 thoracic segments and in A8 (arrows). This ability is partly impaired for *Ubx*<sup>HX</sup> and *Ubx*<sup>UA</sup> with a stronger effect for *Ubx*<sup>UA</sup>. Right panels are enlargement of white boxes.

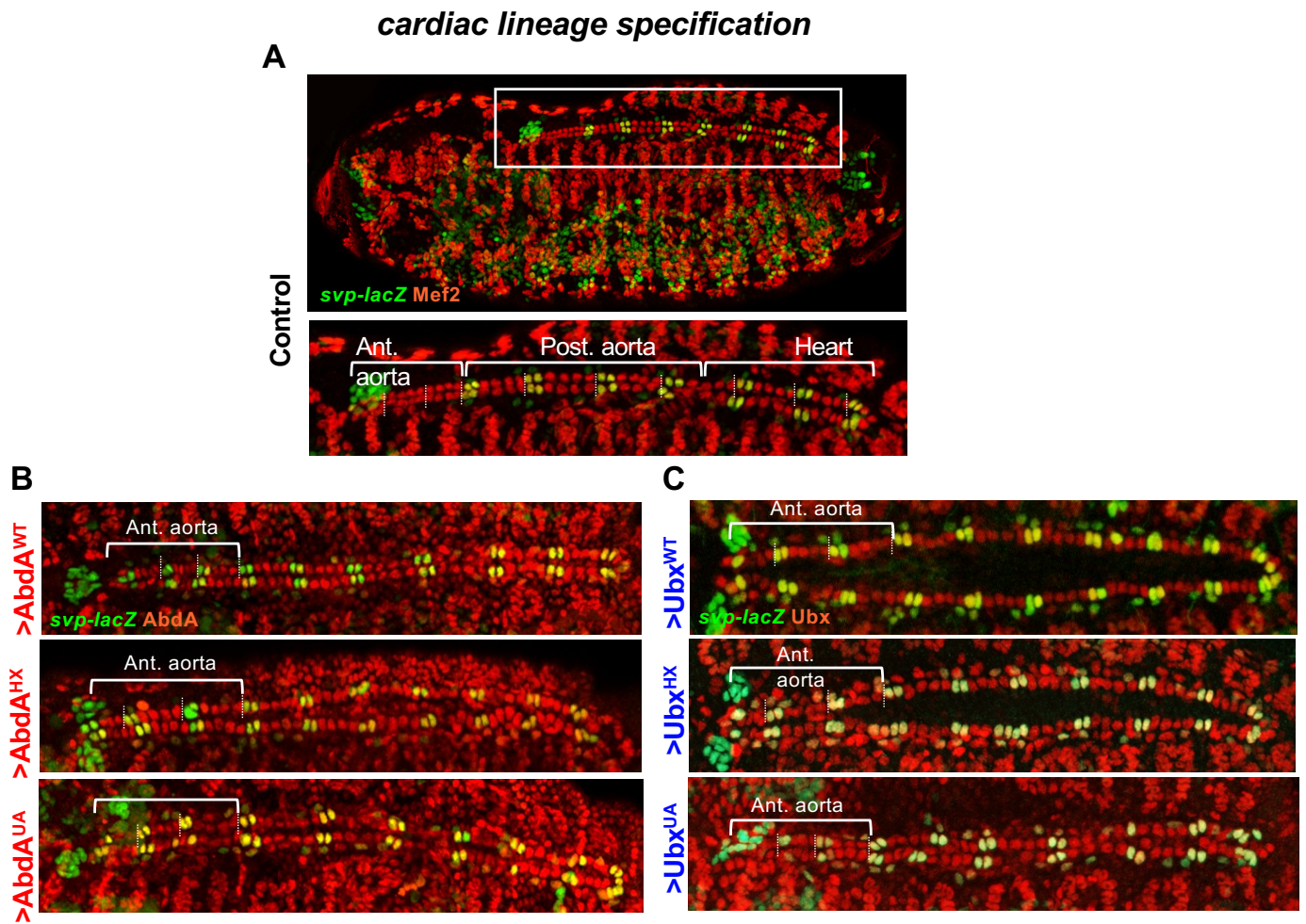

**Figure S3. Ubx and AbdA motif dependency for cardiac cell lineage specification.** **(A)** Control embryo. Axial specialization of the embryonic heart (white box) is illustrated in the magnified view below. Control embryos were stained for Mef2 (red), that marks somatic muscles and cardioblasts, and by *svp* (green). The cardiac lineage is formed by four pairs of cardioblasts per hemi-segment (vertical white dashed lines) in the anterior aorta (ant. aorta) whereas the posterior aorta (post. aorta) and the proper heart contain six pairs of cells, with the two anterior most expressing *svp*. **(B)** *24B-Gal4* driven mesodermal expression of *AbdA<sup>WT</sup>* (red) induces a six-cell lineage in anterior aorta evidenced by *svp* expression. HX and UA mutations do not affect this transformation. **(C)** *24B-Gal4* driven mesodermal expression of *Ubx<sup>WT</sup>* (red) induces a six-cell lineage in anterior aorta while motif mutations partially affect this transformation with a stronger effect seen for the UA compared to the HX mutation.

#### A beard suppression

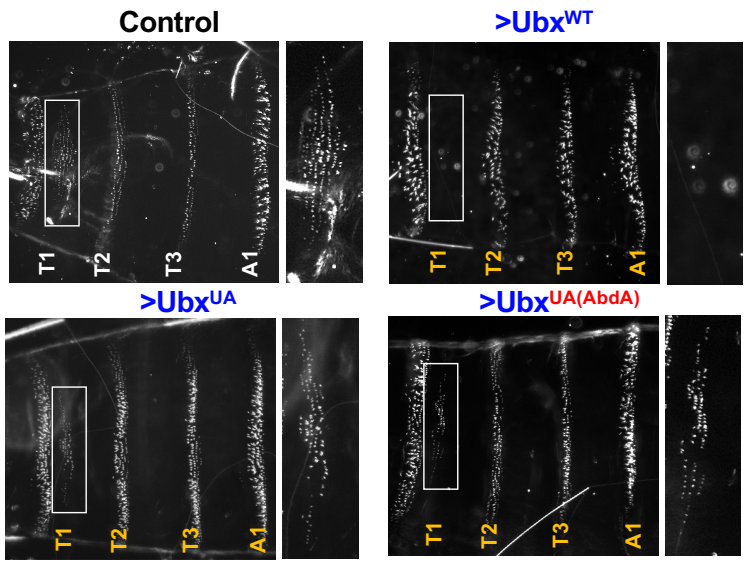

A'

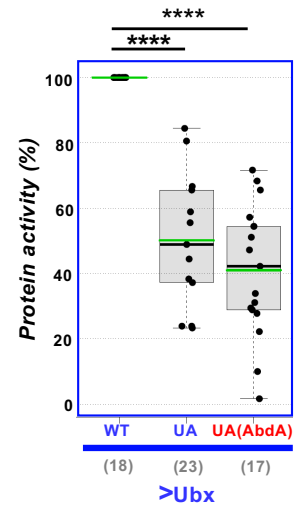

#### B cerebral branch suppression

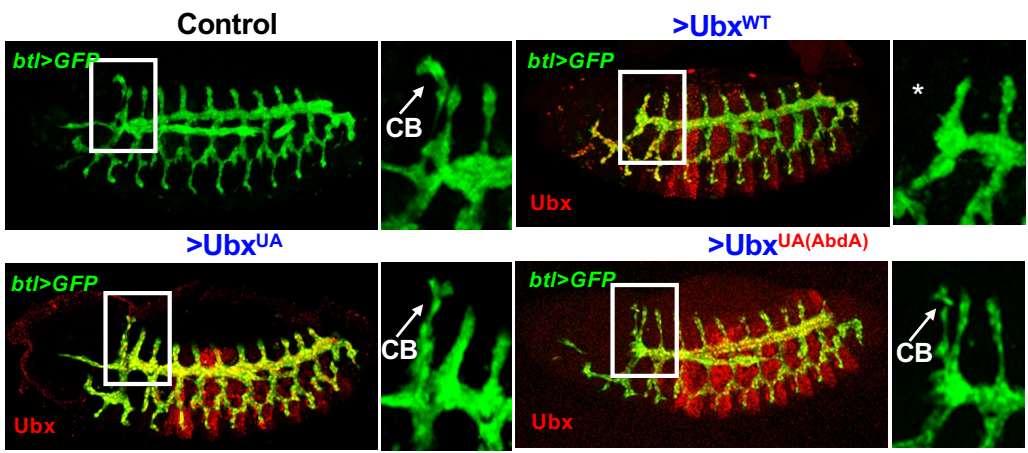

B'

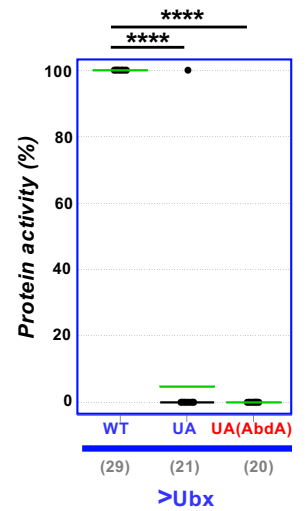

#### C somatic muscle specification

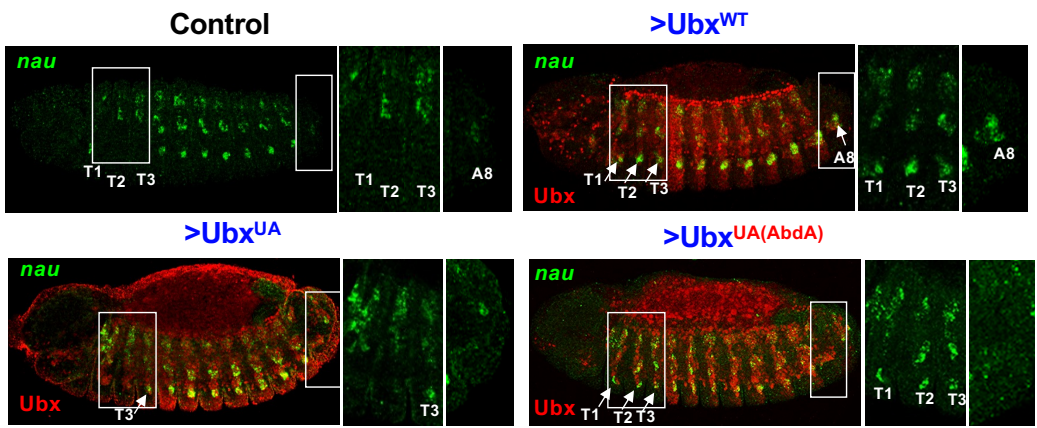

C'

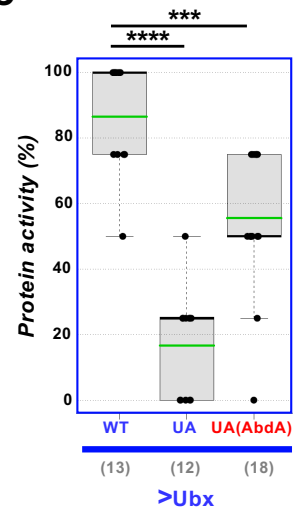

Figure S4 part I

#### D cardiac lineage specification

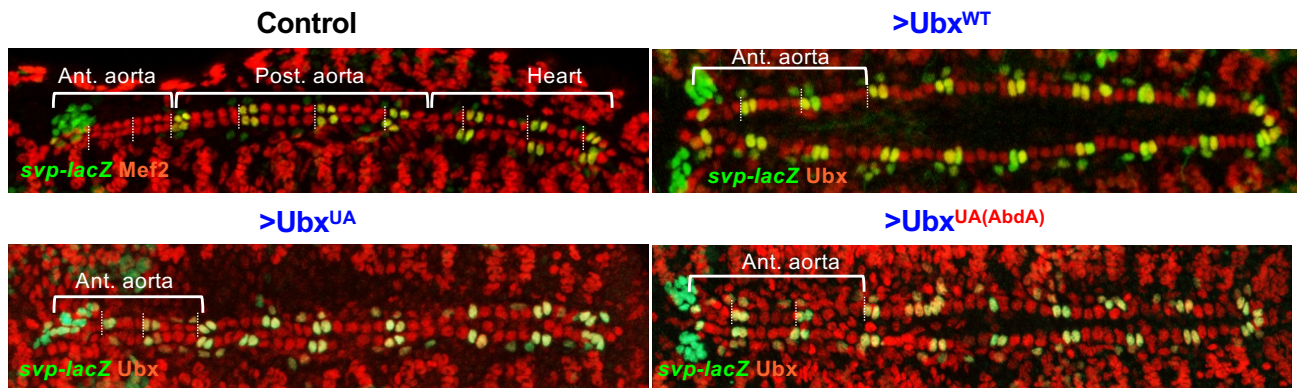

D'

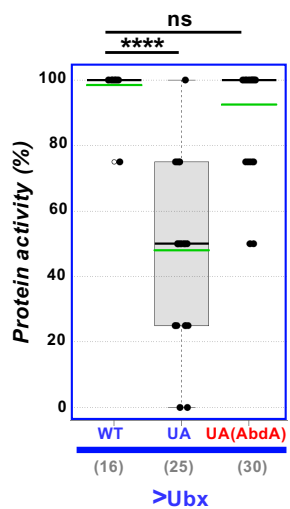

E

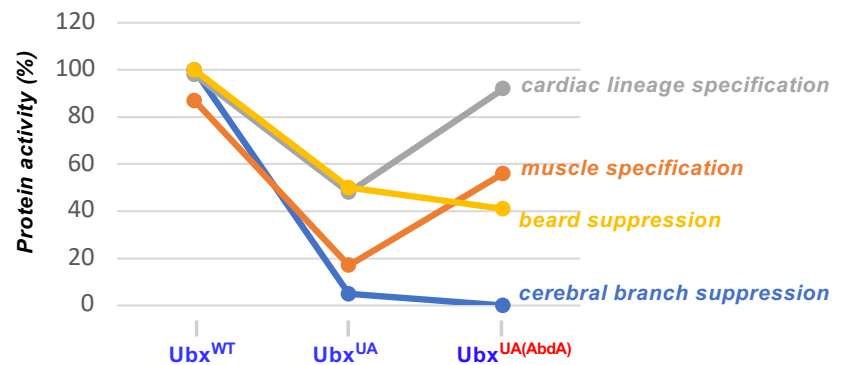

Figure S4 part II

**Figure S4. Protein activity conferred by the AbdA UA motif to Ubx.** (A, A') Beard suppression. (B, B') Cerebral branch suppression. (C, C') Somatic muscle specification. (D, D') Cardiac lineage specification. Ubx<sup>UA(AbdA)</sup> includes the following change: <u>IQA</u>**I**<u>KE</u><u>L</u><u>NE</u><u>Q</u><u>E</u><u>K</u> to <u>LRA</u>**V**<u>K</u><u>E</u><u>I</u><u>N</u><u>E</u><u>Q</u><u>A</u><u>R</u>, which include divergent positions within the UA motif (underlined) and two motif flanking residues (non-bold) changed to AbdA identity to favor proper motif context. Ubx<sup>UA(AbdA)</sup> does not restore activity for beard suppression and cerebral branch repression, while it does so for somatic muscle specification and cardiac lineage specification. Numbers in brackets on graphs indicate the sample size (N) of each experiment. Statistical significance using Fisher or Mann-Whitney tests (see methods) are \*\*\*\* for $p < 0.0001$, \*\*\* for $0.0001 < p < 0.001$ and ns (not significant) for $p > 0.05$. Mean values are in green. (E) Comparison of protein activities of Ubx UA variants.

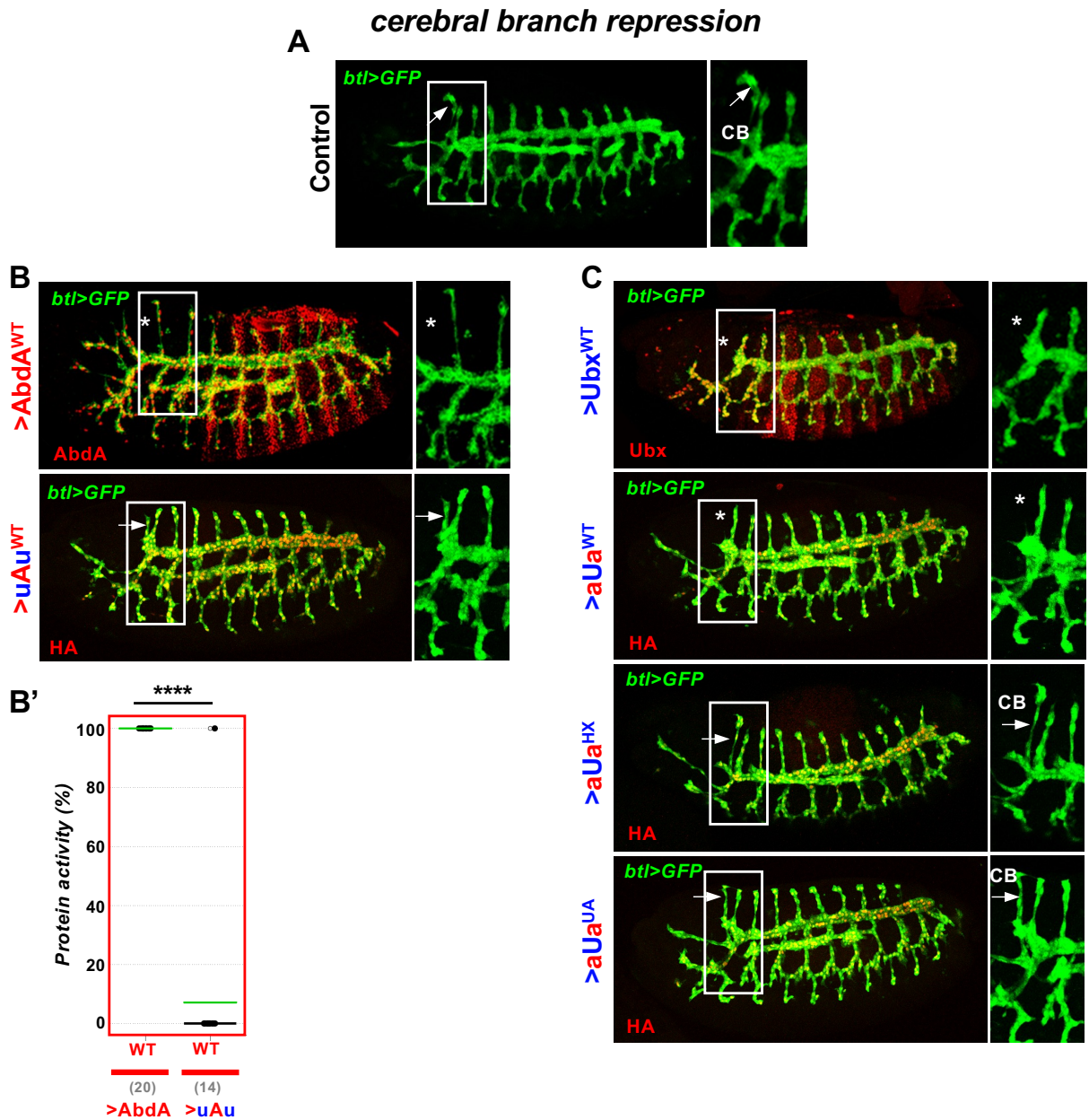

**Figure S5. Motif dependency for uAu and aUa chimeric proteins: cerebral branch suppression.** Trachea are visualized by *btl-Gal4* driven GFP (green). **(A)** In control embryos cerebral branch (CB) forms exclusively in T2 (arrow). Right panels are enlargement of white boxes. **(B, C)** Chimeric protein expression is in red. **(B)** The uAu chimera is almost fully inactive. **(C)** The aUa chimera retains cerebral branch repressive activity (60%). Mutation of the HX or UA motifs within the aUa protein strongly impairs cerebral branch suppression. **(B')** Protein activities quantifications (see methods and Table S2). Numbers in brackets on graphs indicate the sample size (N) of each experiment. Statistical significance using the Fisher test is \*\*\*\* for $p < 0.0001$. Mean values are in green.

#### somatic muscle specification

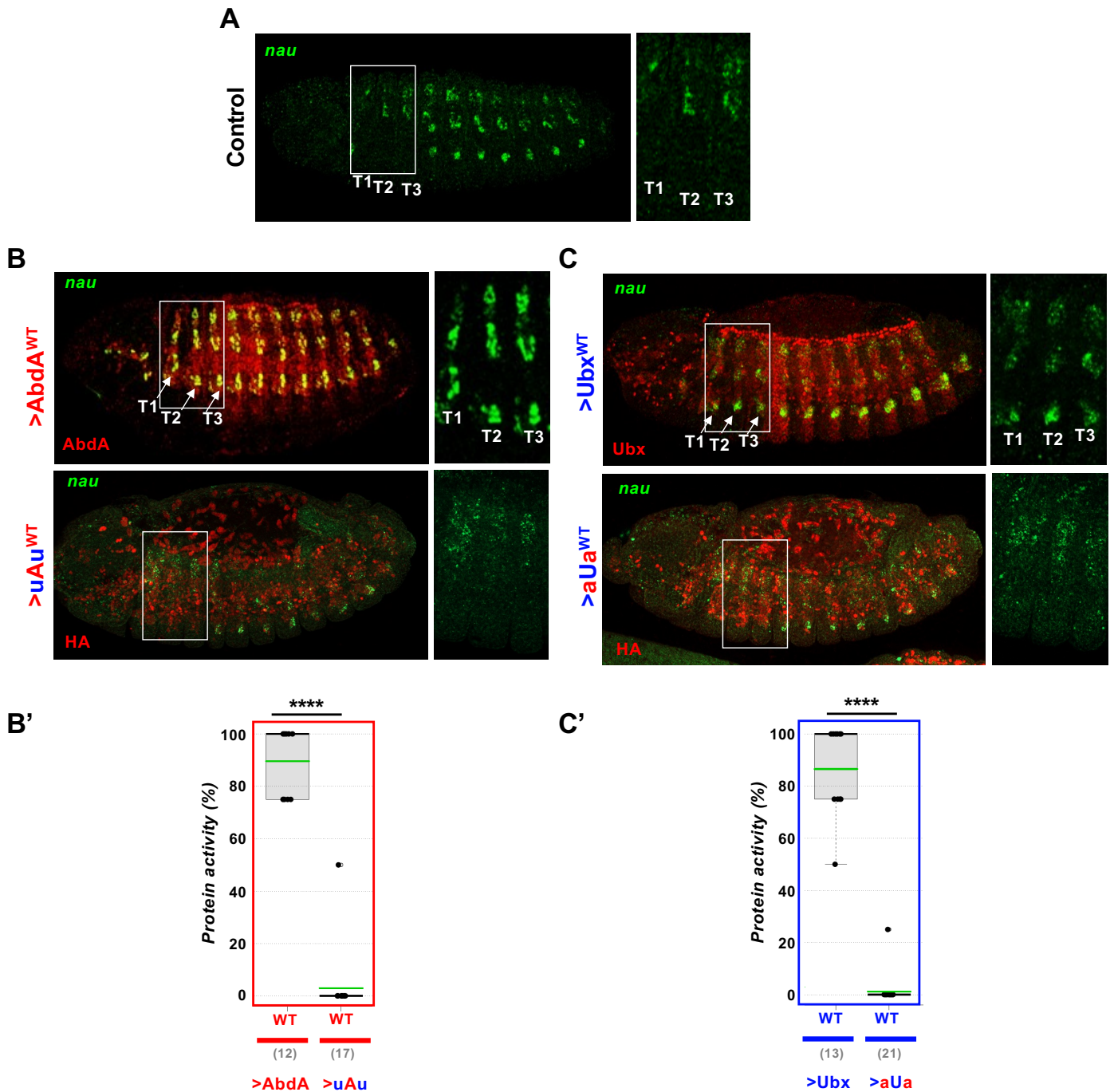

**Figure S6. Motif dependency for uAu and aUa chimeric proteins: somatic muscle specification.** Somatic muscles are visualized by the expression of *nautilus* (*nau*, green). **(A)** control embryo. **(B, C)** Chimeric protein expression (red) was driven by the *24B-Gal4* mesodermal expression. The uAu **(B)** and aUa **(C)** chimeras are inactive. **(B', C')** Protein activity quantifications (see methods and Table S2). Numbers in brackets on graphs indicate the sample size (N) of each experiment. Statistical significance using the Mann-Whitney test is \*\*\*\* for $p < 0.0001$. Mean values are in green.

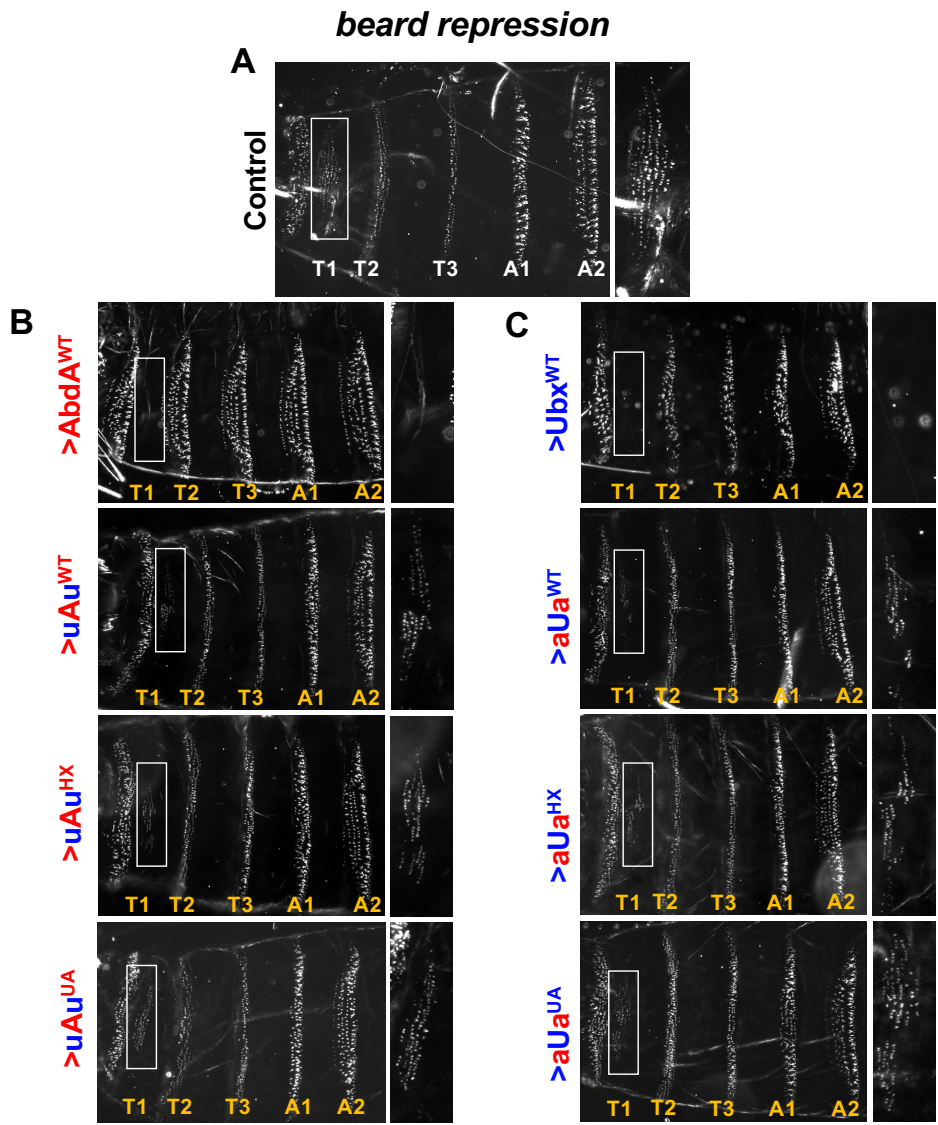

**Figure S7. Motif dependency for uAu and aUa chimeric proteins: beard suppression.** (A) Control. The beard, a group of around 180 small denticles framed in the white rectangle, forms exclusively in the first thoracic segment T1. A magnified view is shown in the right panel. (B, C) Chimeric protein expression was achieved by *arm-Gal4* driven ectodermal expression. Yellow lettering indicates position of segments as referred to WT. The uAu<sup>WT</sup> (B) and aUa<sup>WT</sup> (C) chimeras retain beard suppression activities (52 and 55% respectively). Only mutation of the UA motif within the aUa chimera alters beard suppressive activity.

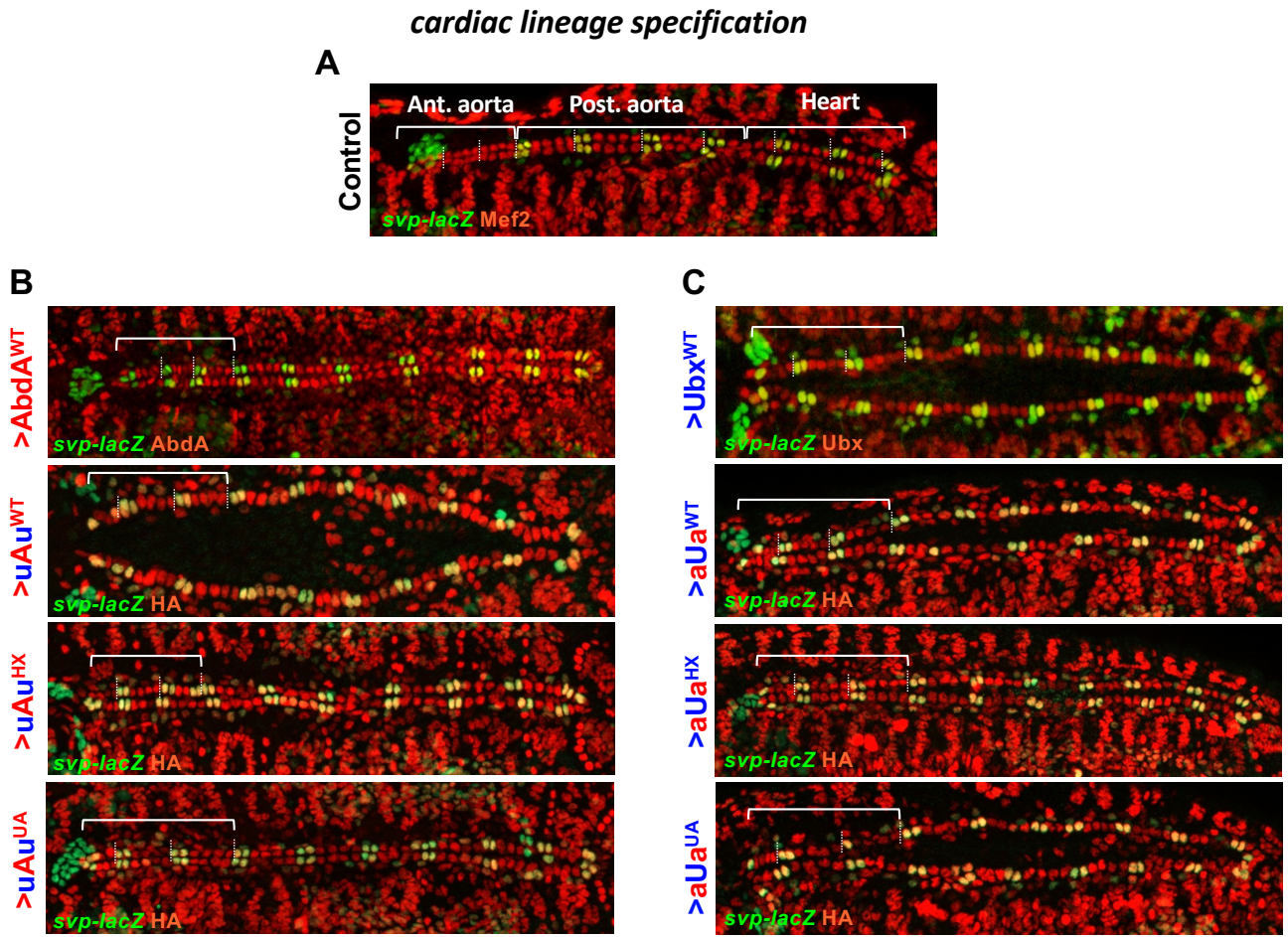

**Figure S8. Motif dependency for uAu and aUa chimeric proteins: cardiac cell lineage specification.** (A) Control embryo stained by Mef2 (red) and *svp* (green). The cardiac lineage is formed by four pairs of cardioblasts per hemi-segment in the anterior aorta (ant. aorta) whereas the posterior aorta (post. aorta) and the proper heart contain six pairs of cells, with the two anterior most expressing *svp*. (B, C) Chimeric protein expression was achieved by *24B-Gal4* driven expression (red). The uAu<sup>WT</sup> (B) and aUa<sup>WT</sup> (C) chimeras retain almost full cardiac cell lineage specification activities. These activities are not affected by HX or UA mutations.

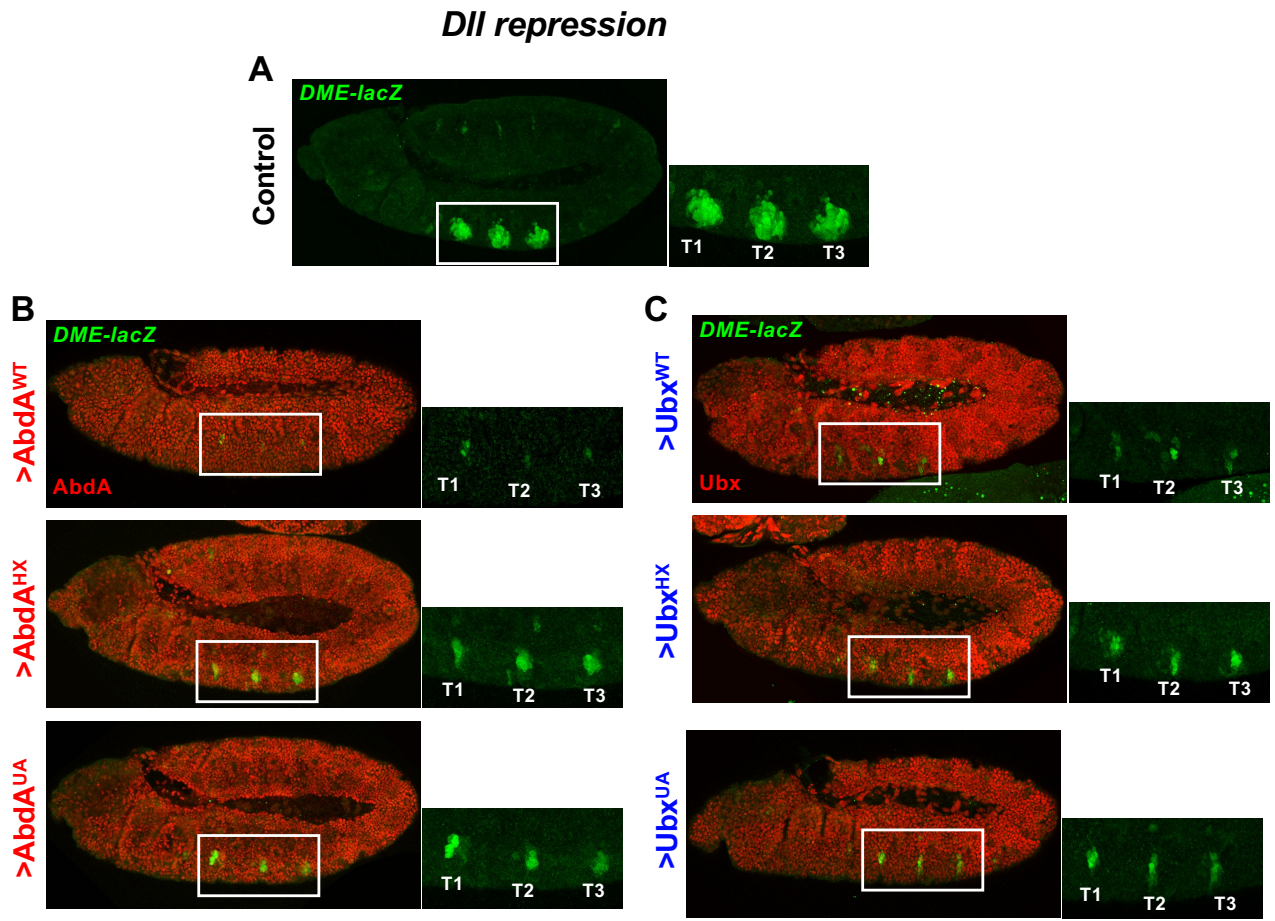

**Figure S9. Ubx and AbdA motif dependency for *Dll* repression suppression.** (A) Control embryo. *Dll* gene expression is confined to the three thoracic segments (T1-T3 in the white rectangle). A magnified view is shown in the right panel. The *DME-lacZ* line recapitulates *Dll* expression and regulation. (B, C) *arm-Gal4* driven ectodermal expression of AbdA (B) or Ubx (C) (red) leads to repression of thoracic *Dll* expression. Repression is diminished upon HX and UA mutations in AbdA, and upon HX mutation in Ubx.

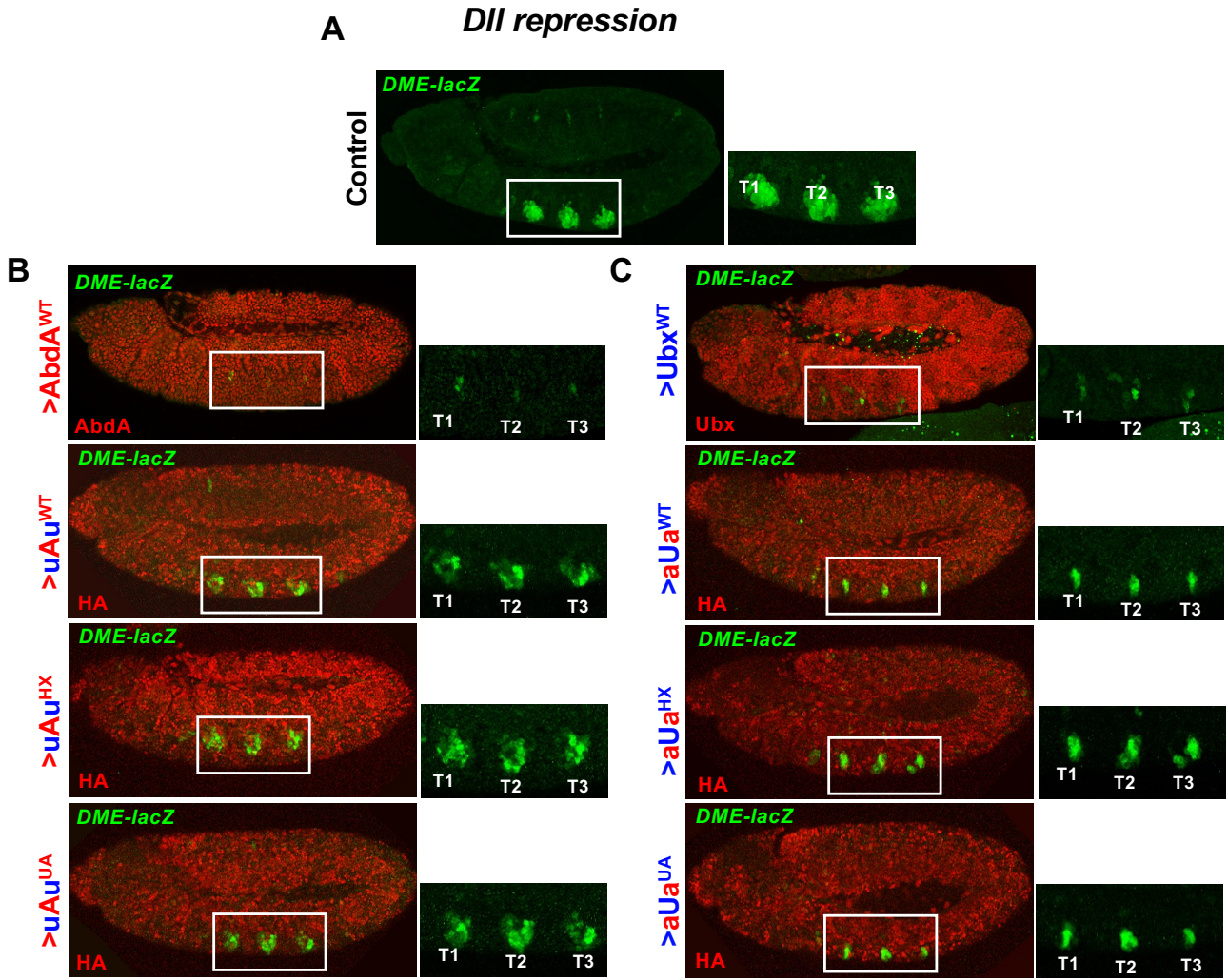

**Figure S10. Motif dependency for uAu and aUa chimeric proteins: *Dll* repression.** (A) Control embryo expressing DME-lacZ reporter gene that recapitulates *Dll* expression and regulation in the three thoracic segments (T1-T3 in the white rectangle and magnified view in the right panel). (B, C) The Chimeric proteins expression was achieved by *arm-Gal4* driven ectodermal expression (red). The uAu<sup>WT</sup> (B) and aUa<sup>WT</sup> (C) chimeras repress *Dll* with different efficiency (50% and 75% respectively). In both cases, full chimera activity requires integrity of the HX motif, but not that of the UA motif.

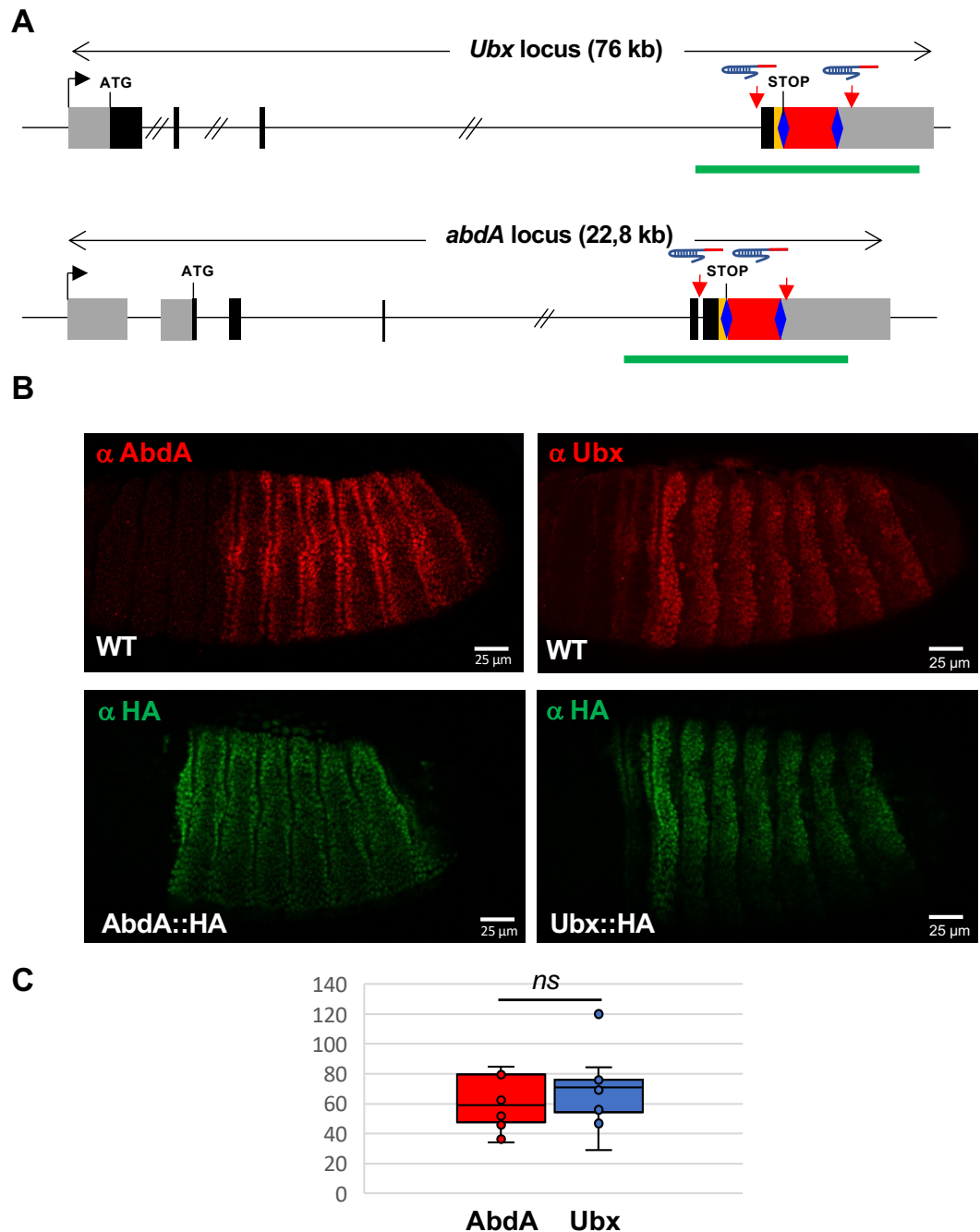

**Figure S11. Compared endogenous Ubx and AbdA expression levels using CRISPR engineered loci.** **(A)** Strategies to generate Ubx and AbdA HA tagged loci. CRISPR modified *Ubx* and *abdA* loci are shown. Coding exons are represented in black, while non-coding exons are in grey. Double slash bars indicate regions that are not to scale. In both cases, the last exon was targeted by homologous recombination to introduce a 3XHA tag (yellow) and a selection cassette (DsRed (red) flanked by LoxP sites (blue lozenges)). The regions targeted by gRNAs (see Table S3 for sequences) are indicated by red arrows. Green lines represent the sequences inserted into the donor constructs for homologous recombination. **(B)** Comparison of expression patterns in wildtype and CRISPR fly lines. Anti-Ubx (mouse Mab FP3.38, DSHB AB\_10805300), anti AbdA (rabbit polyclonal P48, Yacine Graba) and rat anti HA (Roche) anti were used. **(C)** Compared quantification of Ubx::HA and AbdA::HA protein levels. Expression levels vary within their respective patterns: for Ubx expression peaks in the functional domain in pA3/A1 segment, while for AbdA expression is similar in all segments. In addition, both for Ubx and AbdA, expression is higher at segment borders. Quantification was performed within A1 for Ubx, and A2 for AbdA, avoiding the segment borders. Data (see Table S4) show no significant (ns) difference between AbdA::HA and Ubx::HA (p-value of 0,64 in an unpaired t-test).

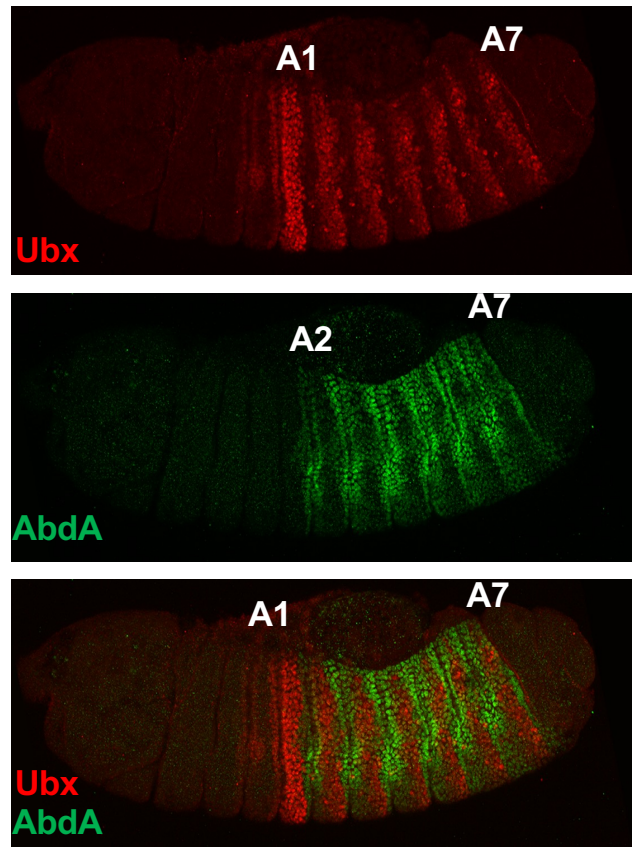

**Figure S12. Overlap of Ubx and AbdA protein expression.**
Images used for Figure 1D. Wild type embryos were stained for Ubx (red) and AbdA (green). A1 (abdominal segment 1); A7 (abdominal segment 7)

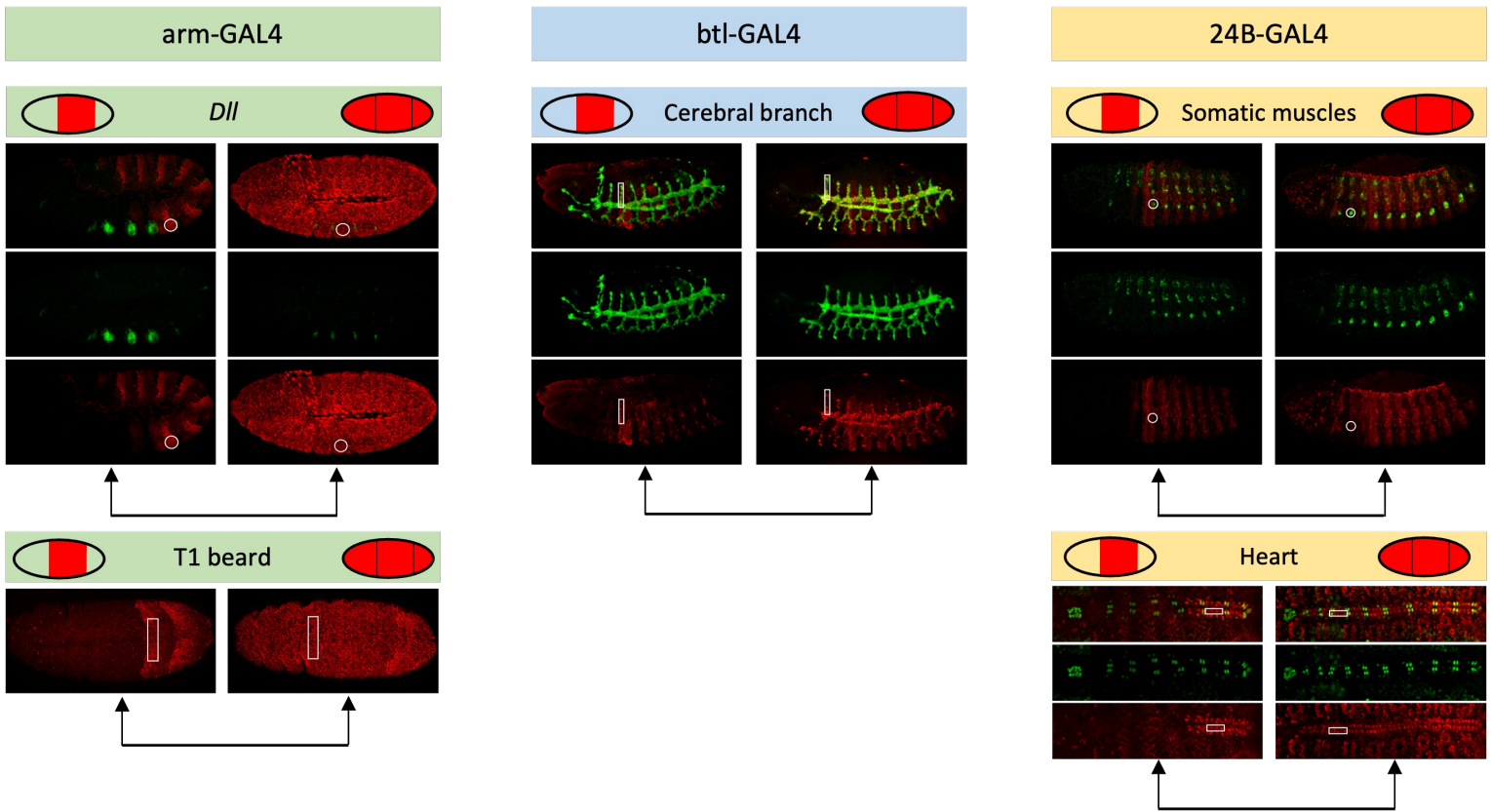

**Figure S13. Procedures for the evaluation of UAS-driven AbdA and Ubx expression levels.** Crosses for the generation of embryos allowing the comparison of endogenous and UAS-driven Ubx and AbdA proteins are depicted on the top of the figure. Regions selected for protein quantification for the different drivers and functional readouts are illustrated below.

|  | Exd-dependency |
| --- | --- |
| Distalless repression | Yes |
| beard repression | No |
| cardiac lineage specification | No |
| somatic muscle specification | No |
| cerebral branch repression | ? |

| **Associated Reference** |
| --- |
| Gebelein, Dev Cell 2002 |
| Sambrani, Developmental dynamics, 2011 |
| Merabet, PLoS Genetics 2011 |
| Michelson, Development 1994 |
| Merabet, EMBO report 2005 |

**MOTIF CONTRIBUTION SUMARY**

Mean of WT prot activity - Mean of motif mutated prot activity

| Protein activity (means, % relative to WT) | Ubx WT | Ubx HX | Ubx UA |
| --- | --- | --- | --- |
| T1 beard | 100 | 41 | 50 |
| cardiac lineage | 98 | 61 | 48 |
| cerebral branch | 100 | 0 | 5 |
| somatic muscle | 87 | 49 | 17 |
| Dll Repression | 78 | 66 | 79 |

| aUa | aUa HX | aUa UA |
| --- | --- | --- |
| 55 | 56 | 33 |
| 88 | 89 | 98 |
| 60 | 12 | 0 |
| 1 | NA | NA |
| 75 | 62 | 73 |

| AbdA WT | AbdA HX | AbdA UA |
| --- | --- | --- |
| 96 | 99 | 90 |
| 93 | 99 | 86 |
| 100 | 100 | 100 |
| 90 | 98 | 94 |
| 95 | 83 | 79 |

| uAu | uAu HX | uAu UA |
| --- | --- | --- |
| 52 | 42 | 41 |
| 89 | 85 | 88 |
| 7 | NA | NA |
| 3 | NA | NA |
| 50 | 38 | 48 |

| Motif contribution (mean of WT prot activity -<br>mean of motif mutated prot activityy) | HX |  | UA |  |
| --- | --- | --- | --- | --- |
|  | Ubx | aUa | Ubx | aUa |
| T1 beard | 59 | -1 | 50 | 22 |
| cardiac lineage | 37 | -1 | 50 | -10 |
| cerebral branch | 100 | 48 | 95 | 60 |
| somatic muscle | 38 | NA | 70 | NA |
| Dll Repression | 12 | 13 | -1 | 2 |

| HX |  | UA |  |
| --- | --- | --- | --- |
| AbdA | uAu | AbdA | uAu |
| -3 | 10 | 6 | 11 |
| -6 | 4 | 7 | 1 |
| 0 | NA | 0 | NA |
| -8 | NA | -4 | NA |
| 12 | 12 | 16 | 2 |

#### T1 Beard

#### Number of beard denticles in T1

| Control | Ubx WT | Ubx HX | Ubx UA | Ubx UA(AbdA) | aUa WT | aUa HX | aUa UA | AbdA WT | AbdA HX | AbdA UA | uAu WT | uAu HX | uAu UA |
| --- | --- | --- | --- | --- | --- | --- | --- | --- | --- | --- | --- | --- | --- |
| 214 | 0 | 59 | 28 | 119 | 46 | 119 | 155 | 24 | 3 | 37 | 115 | 108 | 138 |
| 182 | 0 | 100 | 111 | 53 | 72 | 84 | 164 | 0 | 0 | 24 | 192 | 114 | 149 |
| 106 | 0 | 120 | 137 | 142 | 57 | 74 | 152 | 10 | 0 | 14 | 112 | 12 | 141 |
| 170 | 0 | 107 | 138 | 90 | 127 | 136 | 110 | 0 | 0 | 18 | 61 | 100 | 145 |
| 160 | 0 | 74 | 137 | 129 | 35 | 57 | 107 | 0 | 0 | 7 | 104 | 96 | 98 |
| 222 | 0 | 125 | 80 | 97 | 104 | 18 | 98 | 9 | 0 | 23 | 78 | 105 | 135 |
| 200 | 0 | 120 | 60 | 179 | 104 | 75 | 110 | 0 | 0 | 0 | 28 | 109 | 105 |
| 201 | 0 | 102 | 62 | 132 | 34 | 112 | 86 | 0 | 8 | 20 | 128 | 141 | 32 |
| 170 | 0 | 115 | 35 | 164 | 114 | 90 | 125 | 0 | 0 | 0 | 9 | 104 | 82 |
| 178 | 0 | 107 | 74 | 64 | 165 | 43 | 140 | 3 | 0 | 43 | 52 | 122 | 74 |
| 201 | 0 | 83 | 92 | 130 | 105 | 128 | 156 | 0 | 16 | 0 | 56 | 109 | 134 |
| 178 | 0 | 128 | 100 | 84 | 64 | 60 | 127 | 32 | 0 | 0 | 106 | 104 | 117 |
| 156 | 0 | 106 | 113 | 106 | 42 | 26 | 145 | 0 | 3 | 18 | 96 | 30 | 95 |
| 177 | 0 | 107 |  | 79 | 69 | 28 | 143 | 0 | 0 | 51 | 111 | 113 | 139 |
| 164 | 0 | 136 |  | 126 | 63 | 89 | 125 | 42 | 0 | 31 | 114 | 85 | 15 |
| 182 | 0 | 82 |  | 84 | 90 | 51 | 172 | 0 | 18 | 0 | 79 | 185 |  |
| 152 | 0 | 109 |  | 59 | 75 | 140 | 48 | 7 | 3 | 16 | 19 | 90 |  |
| 193 | 0 | 148 |  |  | 90 | 114 | 95 | 0 | 0 | 10 | 74 | 180 |  |
| 212 |  | 83 |  |  | 104 | 111 | 66 | 0 | 0 | 2 | 58 | 195 |  |
| 212 |  |  |  |  | 114 | 146 | 86 |  |  | 28 | 108 | 97 |  |
| 207 |  |  |  |  | 45 | 70 |  |  |  |  | 111 | 31 |  |
| 125 |  |  |  |  | 52 | 55 |  |  |  |  |  | 64 |  |
|  |  |  |  |  |  | 8 |  |  |  |  |  |  |  |

Mean 180

#### same control applied for all assays

#### Protein activity = Percentage of T1 beard reduction compared to the control

| Ubx WT | Ubx HX | Ubx UA | Ubx UA(AbdA) | aUa WT | aUa HX | aUa UA | AbdA WT | AbdA HX | AbdA UA | uAu WT | uAu HX | uAu UA |
| --- | --- | --- | --- | --- | --- | --- | --- | --- | --- | --- | --- | --- |
| 100 | 67 | 84 | 34 | 74 | 34 | 14 | 87 | 98 | 79 | 36 | 40 | 23 |
| 100 | 44 | 38 | 72 | 60 | 53 | 9 | 100 | 100 | 87 | -7 | 37 | 17 |
| 100 | 33 | 24 | 22 | 68 | 59 | 16 | 94 | 100 | 92 | 38 | 93 | 22 |
| 100 | 41 | 23 | 51 | 29 | 24 | 39 | 100 | 100 | 90 | 66 | 44 | 19 |
| 100 | 59 | 24 | 29 | 81 | 68 | 41 | 100 | 100 | 96 | 42 | 47 | 46 |
| 100 | 31 | 56 | 47 | 42 | 90 | 46 | 95 | 100 | 87 | 57 | 42 | 25 |
| 100 | 33 | 67 | 2 | 42 | 58 | 39 | 100 | 100 | 100 | 84 | 39 | 42 |
| 100 | 43 | 66 | 28 | 81 | 38 | 52 | 100 | 96 | 89 | 29 | 22 | 82 |
| 100 | 36 | 81 | 10 | 37 | 50 | 31 | 100 | 100 | 100 | 95 | 42 | 54 |
| 100 | 41 | 59 | 66 | 8 | 76 | 22 | 98 | 100 | 76 | 71 | 32 | 59 |
| 100 | 54 | 49 | 29 | 42 | 29 | 13 | 100 | 91 | 100 | 69 | 39 | 26 |
| 100 | 29 | 44 | 54 | 64 | 67 | 29 | 82 | 100 | 100 | 41 | 42 | 35 |
| 100 | 41 | 37 | 42 | 77 | 86 | 19 | 100 | 98 | 90 | 47 | 83 | 47 |
| 100 | 41 |  | 57 | 62 | 84 | 21 | 100 | 100 | 72 | 38 | 37 | 23 |
| 100 | 24 |  | 31 | 65 | 51 | 31 | 77 | 100 | 83 | 37 | 53 | 92 |
| 100 | 54 |  | 54 | 50 | 72 | 4 | 100 | 90 | 100 | 56 | -3 |  |
| 100 | 39 |  | 68 | 58 | 22 | 73 | 96 | 98 | 91 | 89 | 50 |  |
| 100 | 18 |  |  | 50 | 37 | 47 | 100 | 100 | 94 | 59 | 0 |  |
|  | 54 |  |  | 42 | 38 | 63 | 100 | 100 | 99 | 68 | -8 |  |
|  |  |  |  | 37 | 19 | 52 |  |  | 84 | 40 | 46 |  |
|  |  |  |  | 75 | 61 |  |  |  |  | 38 | 83 |  |
|  |  |  |  | 71 | 69 |  |  |  |  |  | 64 |  |
|  |  |  |  |  | 96 |  |  |  |  |  |  |  |

|  | Ubx WT | Ubx HX | Ubx UA | Ubx UA(AbdA) |  | aUa WT | aUa HX | aUa UA |  | AbdA WT | AbdA HX | AbdA UA |  | uAu WT | uAu HX | uAu UA |
| --- | --- | --- | --- | --- | --- | --- | --- | --- | --- | --- | --- | --- | --- | --- | --- | --- |
| n | 18 | 19 | 13 | 17 |  | 22 | 23 | 20 |  | 19 | 19 | 20 |  | 21 | 22 | 15 |
| Mean | 100 | 41 | 50 | 41 |  | 55 | 56 | 33 |  | 96 | 99 | 90 |  | 52 | 42 | 41 |
| SD | 0 | 12 | 21 | 20 |  | 19 | 23 | 19 |  | 7 | 3 | 9 |  | 23 | 25 | 23 |

|  | HX to Ubx | UA to Ubx | UA(AbdA) to Ubx | HX to aUa | UA to aUa | HX to AbdA | UA to AbdA | HX to uAu | UA to uAu |
| --- | --- | --- | --- | --- | --- | --- | --- | --- | --- |
| Motif contribution* | 59 | 50 | 59 | -1 | 22 | -3 | 6 | 10 | 11 |

\*Motif contribution = Mean of WT prot activity - Mean of motif mutated prot activity

##### Cerebral Branch

Cerebral branch absent = 0 ; Cerebral branch present =1

[illegible]

Protein activity: Cerebral branch absent = 100 % ; Cerebral branch present = 0%

[illegible]

|  | Ubx WT | Ubx HX | Ubx UA | Ubx UA(AbΔa) | aUA WT | aUA HX | aUA UA | AbΔa WT | AbΔa HX | AbΔa UA | uAU WT | uAU HX | uAU UA |
| --- | --- | --- | --- | --- | --- | --- | --- | --- | --- | --- | --- | --- | --- |
| n | 29 | 20 | 21 | 20 | 20 | 26 | 14 | 20 | 20 | 20 | 14 | NA | NA |
| Mean | 100 | 0 | 5 | 0 | 60 | 12 | 0 | 100 | 100 | 100 | 7 |  |  |
| SD | 0 | 0 | 22 | 0 | 50 | 33 | 0 | 0 | 0 | 0 | 27 |  |  |

|  | HX to Ubx |  |  | UA to Ubx |  | UA(AbdA) to Ubx |  | HX to aUa |  | UA to aUa |  | HX to AbdA |  | UA to AbdA |  | HX to uAu |  | UA to uAu |
| --- | --- | --- | --- | --- | --- | --- | --- | --- | --- | --- | --- | --- | --- | --- | --- | --- | --- | --- |
| Motif contribution* | 100 | 95 | 100 | 48 | 60 |  |  | 0 | 0 |  |  | NA | NA |  |  | NA | NA |  |

\*Motif contribution = Mean of WT prot activity - Mean of motif mutated prot activity

#### Cardiac Lineage

Number of cardiac cells counted in the two posterior most segments of the anterior aorta

| Ubx WT | Ubx HX | Ubx UA | Ubx UA(AbdA) | aUa WT | aUa HX | aUa UA | AbdA WT | AbdA HX | AbdA UA | uAu WT | uAu HX | uAu UA |
| --- | --- | --- | --- | --- | --- | --- | --- | --- | --- | --- | --- | --- |
| 12 | 12 | 10 | 12 | 11 | 12 | 12 | 12 | 12 | 12 | 9 | 12 | 12 |
| 12 | 9 | 12 | 12 | 12 | 10 | 12 | 12 | 12 | 12 | 12 | 12 | 12 |
| 12 | 12 | 10 | 12 | 11 | 10 | 12 | 12 | 12 | 12 | 12 | 11 | 12 |
| 12 | 11 | 10 | 12 | 12 | 12 | 12 | 12 | 11 | 12 | 12 | 12 | 12 |
| 12 | 10 | 9 | 12 | 11 | 12 | 12 | 12 | 12 | 12 | 12 | 12 | 12 |
| 12 | 10 | 11 | 12 | 11 | 10 | 12 | 12 | 12 | 12 | 12 | 10 | 9 |
| 12 | 11 | 10 | 12 | 12 | 12 | 12 | 12 | 12 | 12 | 11 | 12 | 12 |
| 12 | 10 | 10 | 12 | 12 | 12 | 12 | 12 | 12 | 12 | 12 | 11 | 12 |
| 12 | 11 | 10 | 12 | 12 | 12 | 11 | 12 | 12 | 11 | 12 | 11 | 12 |
| 12 | 10 | 8 | 11 | 12 | 12 | 12 | 12 | 12 | 12 | 12 | 11 | 12 |
| 11 | 10 | 11 | 12 | 12 | 12 | 12 | 12 | 12 | 9 | 12 | 12 | 12 |
| 12 | 12 | 9 | 12 | 12 | 12 | 11 | 12 | 12 | 11 | 11 | 12 | 12 |
| 12 | 8 | 9 | 10 | 12 | 11 | 12 | 11 | 12 | 12 | 11 | 11 | 12 |
| 12 | 12 | 10 | 11 | 9 | 12 | 12 | 9 | 12 | 10 | 12 | 11 | 12 |
| 12 | 12 | 11 | 12 | 10 | 12 | 12 | 11 | 12 | 12 | 10 | 11 | 11 |
| 12 | 9 | 10 | 11 | 12 | 11 | 12 | 12 | 12 | 12 | 12 | 11 | 11 |
| 10 | 10 | 11 | 11 | 12 | 11 | 12 | 12 | 12 | 9 | 12 | 12 | 12 |
| 9 | 10 | 12 |  | 12 | 12 | 12 | 12 | 12 | 12 |  |  | 12 |
| 11 | 11 | 12 |  |  | 12 | 12 |  |  |  |  |  | 11 |
| 9 | 10 | 12 |  |  | 12 | 12 |  |  |  |  |  | 12 |
| 10 | 9 | 12 |  |  |  |  |  |  |  |  |  | 10 |
| 12 | 9 | 12 |  |  |  |  |  |  |  |  |  | 12 |
| 10 | 8 | 12 |  |  |  |  |  |  |  |  |  | 9 |
|  | 11 | 12 |  |  |  |  |  |  |  |  |  | 11 |
|  | 9 | 12 |  |  |  |  |  |  |  |  |  | 12 |
|  |  | 12 |  |  |  |  |  |  |  |  |  | 12 |
|  |  | 11 |  |  |  |  |  |  |  |  |  |  |
|  |  | 12 |  |  |  |  |  |  |  |  |  |  |
|  |  | 12 |  |  |  |  |  |  |  |  |  |  |
|  |  | 10 |  |  |  |  |  |  |  |  |  |  |

Protein activity: 12 cardiac cells = 100% ; 11 cardiac cells = 75% ; 10 cardiac cells = 50% ; 9 cardiac cells = 25% ; 8 cardiac cells = 0%

| Ubx WT | Ubx HX | Ubx UA | Ubx UA(AbdA) | aUa WT | aUa HX | aUa UA | AbdA WT | AbdA HX | AbdA UA | uAu WT | uAu HX | uAu UA |
| --- | --- | --- | --- | --- | --- | --- | --- | --- | --- | --- | --- | --- |
| 100 | 100 | 50 | 100 | 75 | 100 | 100 | 100 | 100 | 100 | 25 | 100 | 100 |
| 100 | 25 | 100 | 100 | 100 | 50 | 100 | 100 | 100 | 100 | 100 | 100 | 100 |
| 100 | 100 | 50 | 100 | 75 | 50 | 100 | 100 | 100 | 100 | 100 | 75 | 100 |
| 100 | 75 | 50 | 100 | 100 | 100 | 100 | 100 | 75 | 100 | 100 | 100 | 100 |
| 100 | 50 | 25 | 100 | 75 | 100 | 100 | 100 | 100 | 100 | 100 | 100 | 100 |
| 100 | 50 | 75 | 100 | 75 | 50 | 100 | 100 | 100 | 100 | 100 | 50 | 25 |
| 100 | 75 | 50 | 100 | 100 | 100 | 100 | 100 | 100 | 100 | 75 | 100 | 100 |
| 100 | 50 | 50 | 100 | 100 | 100 | 100 | 100 | 100 | 100 | 100 | 75 | 100 |
| 100 | 75 | 50 | 100 | 100 | 100 | 75 | 100 | 100 | 75 | 100 | 75 | 100 |
| 100 | 50 | 0 | 75 | 100 | 100 | 100 | 100 | 100 | 100 | 100 | 75 | 100 |
| 75 | 50 | 75 | 100 | 100 | 100 | 100 | 100 | 100 | 25 | 100 | 100 | 100 |
| 100 | 100 | 25 | 100 | 100 | 100 | 75 | 100 | 100 | 75 | 100 | 100 | 100 |
| 100 | 0 | 25 | 50 | 100 | 75 | 100 | 75 | 100 | 100 | 75 | 75 | 100 |
| 100 | 100 | 50 | 75 | 25 | 100 | 100 | 25 | 100 | 50 | 100 | 75 | 100 |
| 100 | 100 | 75 | 100 | 50 | 100 | 100 | 75 | 100 | 100 | 50 | 75 | 75 |
| 100 | 25 | 50 | 75 | 100 | 75 | 100 | 100 | 100 | 100 | 100 | 75 | 75 |
| 50 | 50 | 75 | 75 | 100 | 75 | 100 | 100 | 100 | 25 | 100 | 100 | 100 |
| 25 | 50 | 100 | 100 | 100 | 100 | 100 | 100 | 100 | 100 | 100 |  | 100 |
| 75 | 75 | 100 | 100 |  | 100 | 100 |  | 100 |  |  |  | 75 |
| 25 | 50 | 100 | 100 |  | 100 | 100 |  |  |  |  |  | 100 |
| 50 | 25 | 100 | 100 |  |  |  |  |  |  |  |  | 50 |
| 100 | 25 | 100 | 100 |  |  |  |  |  |  |  |  | 100 |
| 50 | 0 | 100 | 100 |  |  |  |  |  |  |  |  | 25 |
|  | 75 | 100 | 100 |  |  |  |  |  |  |  |  | 75 |
|  | 25 | 100 | 100 |  |  |  |  |  |  |  |  | 100 |
|  |  | 100 | 100 |  |  |  |  |  |  |  |  |  |
|  |  | 75 | 100 |  |  |  |  |  |  |  |  |  |
|  |  | 100 | 100 |  |  |  |  |  |  |  |  |  |
|  |  | 100 | 100 |  |  |  |  |  |  |  |  |  |
|  |  | 50 |  |  |  |  |  |  |  |  |  |  |

|  | Ubx WT | Ubx HX | Ubx UA | Ubx UA(AbdA) | aUa WT | aUa HX | aUa UA | AbdA WT | AbdA HX | AbdA UA | uAu WT | uAu HX | uAu UA |
| --- | --- | --- | --- | --- | --- | --- | --- | --- | --- | --- | --- | --- | --- |
| n | 16 | 23 | 25 | 30 | 18 | 20 | 20 | 18 | 19 | 18 | 18 | 17 | 26 |
| Mean | 98 | 61 | 48 | 93 | 88 | 89 | 98 | 93 | 99 | 86 | 89 | 85 | 88 |
| SD | 6 | 30 | 25 | 15 | 21 | 19 | 8 | 19 | 6 | 26 | 21 | 15 | 23 |

| Motif contribution* | HX to Ubx | UA to Ubx | UA(AbdA) to Ubx | HX to aUa | UA to aUa | HX to AbdA | UA to AbdA | HX to uAu | UA to uAu |
| --- | --- | --- | --- | --- | --- | --- | --- | --- | --- |
|  | 37 | 50 | 5 | -1 | -10 | -6 | 7 | 4 | 1 |

\*Motif contribution = Mean of WT prot activity - Mean of motif mutated prot activity

Somatic Muscle

Number of nautilus additional ventral cluster induced per embryo

| Ubx WT | UbxHX | UbxUA | Ubx UA(AbdA) | aUa WT | aUa HX | aUa UA | AbdA WT | AbdA HX | AbdA UA | uAu WT | uAu HX | uAu UA |
| --- | --- | --- | --- | --- | --- | --- | --- | --- | --- | --- | --- | --- |
| 4 | 3 | 2 | 3 | 1 | NA | NA | 4 | 4 | 4 | 2 | NA | NA |
| 4 | 3 | 1 | 3 | 0 |  |  | 4 | 4 | 4 | 0 |  |  |
| 4 | 3 | 1 | 3 | 0 |  |  | 4 | 4 | 4 | 0 |  |  |
| 4 | 3 | 1 | 3 | 0 |  |  | 4 | 4 | 4 | 0 |  |  |
| 4 | 3 | 1 | 3 | 0 |  |  | 4 | 4 | 4 | 0 |  |  |
| 4 | 3 | 1 | 3 | 0 |  |  | 4 | 4 | 4 | 0 |  |  |
| 4 | 2 | 1 | 3 | 0 |  |  | 4 | 4 | 4 | 0 |  |  |
| 3 | 2 | 0 | 2 | 0 |  |  | 3 | 4 | 4 | 0 |  |  |
| 3 | 2 | 0 | 2 | 0 |  |  | 3 | 4 | 3 | 0 |  |  |
| 3 | 2 | 0 | 2 | 0 |  |  | 3 | 4 | 3 | 0 |  |  |
| 3 | 2 | 0 | 2 | 0 |  |  | 3 | 3 | 3 | 0 |  |  |
| 2 | 2 |  | 2 | 0 |  |  |  |  |  | 0 |  |  |
|  | 2 |  | 2 | 0 |  |  |  |  |  | 0 |  |  |
|  | 1 |  | 2 | 0 |  |  |  |  |  | 0 |  |  |
|  | 0 |  | 2 | 0 |  |  |  |  |  | 0 |  |  |
|  | 0 |  | 1 | 0 |  |  |  |  |  | 0 |  |  |
|  | 0 |  | 0 | 0 |  |  |  |  |  | 0 |  |  |
|  |  |  |  | 0 |  |  |  |  |  |  |  |  |
|  |  |  |  | 0 |  |  |  |  |  |  |  |  |
|  |  |  |  | 0 |  |  |  |  |  |  |  |  |

Protein activity: 4 additional clusters = 100% ; 3 additional clusters = 75% ; 2 additional clusters = 50% ; 1 additional cluster = 25% ; 0 additional cluster = 0%

| Ubx WT | UbxHX | UbxUA | Ubx UA(AbdA) | aUa WT | aUa HX | aUa UA | AbdA WT | AbdA HX | AbdA UA | uAu WT | uAu HX | uAu UA |
| --- | --- | --- | --- | --- | --- | --- | --- | --- | --- | --- | --- | --- |
| 100 | 75 | 50 | 75 | 25 | NA | NA | 100 | 100 | 100 | 50 | NA | NA |
| 100 | 75 | 25 | 75 | 0 |  |  | 100 | 100 | 100 | 0 |  |  |
| 100 | 75 | 25 | 75 | 0 |  |  | 100 | 100 | 100 | 0 |  |  |
| 100 | 75 | 25 | 75 | 0 |  |  | 100 | 100 | 100 | 0 |  |  |
| 100 | 75 | 25 | 75 | 0 |  |  | 100 | 100 | 100 | 0 |  |  |
| 100 | 75 | 25 | 75 | 0 |  |  | 100 | 100 | 100 | 0 |  |  |
| 100 | 50 | 25 | 75 | 0 |  |  | 100 | 100 | 100 | 0 |  |  |
| 75 | 50 | 0 | 50 | 0 |  |  | 75 | 100 | 100 | 0 |  |  |
| 75 | 50 | 0 | 50 | 0 |  |  | 75 | 100 | 100 | 0 |  |  |
| 75 | 50 | 0 | 50 | 0 |  |  | 75 | 100 | 75 | 0 |  |  |
| 75 | 50 | 0 | 50 | 0 |  |  | 75 | 100 | 75 | 0 |  |  |
| 75 | 50 | 0 | 50 | 0 |  |  | 75 | 75 | 75 | 0 |  |  |
| 50 | 50 |  | 50 | 0 |  |  |  |  |  | 0 |  |  |
|  | 50 |  | 50 | 0 |  |  |  |  |  | 0 |  |  |
|  | 25 |  | 50 | 0 |  |  |  |  |  | 0 |  |  |
|  | 0 |  | 50 | 0 |  |  |  |  |  | 0 |  |  |
|  | 0 |  | 25 | 0 |  |  |  |  |  | 0 |  |  |
|  | 0 |  | 0 | 0 |  |  |  |  |  |  |  |  |
|  |  |  |  | 0 |  |  |  |  |  |  |  |  |
|  |  |  |  | 0 |  |  |  |  |  |  |  |  |

|  | Ubx WT | UbxHX | UbxUA | Ubx UA(AbdA) | aUa WT | aUa HX | aUa UA | AbdA WT | AbdA HX | AbdA UA | uAu WT | uAu HX | uAu UA |
| --- | --- | --- | --- | --- | --- | --- | --- | --- | --- | --- | --- | --- | --- |
| n | 13 | 18 | 12 | 18 | 21 | NA | NA | 12 | 12 | 12 | 17 | NA | NA |
| Mean | 87 | 49 | 17 | 56 | 1 |  |  | 90 | 98 | 94 | 3 |  |  |
| SD | 17 | 26 | 16 | 20 | 5 |  |  | 13 | 7 | 11 | 12 |  |  |

| Motif contribution* | HX to Ubx | UA to Ubx | UA(AbdA) to Ubx | HX to aUa | UA to aUa | HX to AbdA | UA to AbdA | HX to uAu | UA to uAu |
| --- | --- | --- | --- | --- | --- | --- | --- | --- | --- |
|  | 38 | 70 | 31 | NA | NA | -8 | -4 | NA | NA |

\*Motif contribution = Mean of WT prot activity - Mean of motif mutated prot activity

#### DII

Mean of the fluorescent DME-lacZ cells area measured in the three thoracic segments

| Control | Ubx WT | Ubx HX | Ubx UA | Control | aUa WT | aUa HX | aUa UA | Control | AbdA WT | AbdA HX | AbdA UA | Control | uAu WT | uAu HX | uAu UA |
| --- | --- | --- | --- | --- | --- | --- | --- | --- | --- | --- | --- | --- | --- | --- | --- |
| 810 | 96 | 229 | 210 | 880 | 123 | 436 | 237 | 810 | 11 | 176 | 327 | 880 | 523 | 657 | 524 |
| 884 | 106 | 333 | 84 | 810 | 140 | 346 | 210 | 884 | 13 | 192 | 231 | 810 | 428 | 674 | 284 |
| 817 | 106 | 245 | 136 | 1104 | 264 | 280 | 225 | 817 | 78 | 212 | 89 | 1104 | 446 | 406 | 405 |
| 933 | 229 | 274 | 187 | 836 | 329 | 293 | 266 | 933 | 20 | 127 | 84 | 836 | 567 | 458 | 345 |
| 919 | 188 | 224 | 152 | 823 | 154 | 427 | 207 | 919 | 79 | 105 | 83 | 823 | 683 | 619 | 288 |
| 519 | 192 | 307 | 189 | 814 | 181 | 350 | 267 | 519 | 44 | 60 | 112 | 814 | 316 | 547 | 476 |
| 671 | 231 | 257 | 231 | 828 | 157 | 336 | 241 | 671 | 44 | 82 | 220 | 828 | 549 | 321 | 495 |
| 863 | 105 | 416 | 149 | 950 | 264 | 392 | 377 | 863 | 35 | 163 | 151 | 950 | 297 | 528 | 422 |
| 642 | 199 | 209 |  | 818 | 295 | 320 | 228 | 642 | 6 | 78 | 159 | 818 | 247 | 552 | 453 |
|  | 193 | 202 |  | 809 | 212 | 395 | 287 |  | 39 | 150 |  | 809 | 406 | 557 | 447 |
|  | 216 | 236 |  | 933 | 208 | 285 | 194 |  |  |  |  | 933 | 283 | 515 | 387 |
|  | 168 |  |  | 890 | 309 | 332 | 172 |  |  |  |  | 890 | 700 | 729 | 312 |
|  | 171 |  |  |  | 231 | 318 | 187 |  |  |  |  |  | 398 | 485 | 314 |
|  | 197 |  |  |  | 193 | 222 | 195 |  |  |  |  |  | 345 | 581 | 544 |
|  |  |  |  |  |  | 192 |  |  |  |  |  |  | 398 | 521 | 624 |
|  |  |  |  |  |  |  |  |  |  |  |  |  |  |  | 721 |
|  |  |  |  |  |  |  |  |  |  |  |  |  |  |  | 608 |
|  |  |  |  |  |  |  |  |  |  |  |  |  |  |  | 599 |
| mean=784 |  |  |  | mean=874 |  |  |  | mean=784 |  |  |  | mean=874 |  |  |  |

Protein activity = Percentage of fluorescent DME-lacZ cells area reduction by comparison to the control.

|  |  |  |  |  |  |  |  |  |  |  |  |  |
| --- | --- | --- | --- | --- | --- | --- | --- | --- | --- | --- | --- | --- |
|  | Ub<sub>x</sub> WT | Ub<sub>x</sub> HX | Ub<sub>x</sub> UA | aUa WT | aUa HX | aUa UA | AbdA WT | AbdA HX | AbdA UA | uAu WT | uAu HX | uAu UA |
|  | 88 | 71 | 73 | 86 | 50 | 73 | 99 | 78 | 58 | 40 | 25 | 40 |
|  | 87 | 57 | 89 | 84 | 60 | 76 | 98 | 76 | 71 | 51 | 23 | 68 |
|  | 87 | 69 | 83 | 70 | 68 | 74 | 90 | 73 | 89 | 49 | 54 | 54 |
|  | 71 | 65 | 76 | 62 | 66 | 70 | 97 | 84 | 89 | 35 | 48 | 61 |
|  | 76 | 71 | 81 | 82 | 51 | 76 | 90 | 87 | 89 | 22 | 29 | 67 |
|  | 76 | 61 | 76 | 79 | 60 | 70 | 94 | 92 | 86 | 64 | 37 | 46 |
|  | 70 | 67 | 71 | 82 | 62 | 72 | 94 | 90 | 72 | 37 | 63 | 43 |
|  | 87 | 47 | 81 | 70 | 55 | 57 | 96 | 79 | 81 | 66 | 40 | 52 |
|  | 75 | 73 |  | 66 | 63 | 74 | 99 | 90 | 80 | 72 | 37 | 48 |
|  | 75 | 74 |  | 76 | 55 | 67 | 95 | 81 |  | 54 | 36 | 49 |
|  | 73 | 70 |  | 76 | 67 | 78 |  |  |  | 68 | 41 | 56 |
|  | 79 |  |  | 65 | 62 | 80 |  |  |  | 20 | 17 | 64 |
|  | 78 |  |  | 74 | 64 | 79 |  |  |  | 54 | 45 | 64 |
|  | 75 |  |  | 78 | 75 | 78 |  |  |  | 61 | 34 | 38 |
|  |  |  |  |  | 78 |  |  |  |  | 54 | 40 | 29 |
|  |  |  |  |  |  |  |  |  |  |  |  | 18 |
|  |  |  |  |  |  |  |  |  |  |  |  | 31 |
|  |  |  |  |  |  |  |  |  |  |  |  | 32 |
|  | Ub<sub>x</sub> WT | Ub<sub>x</sub> HX | Ub<sub>x</sub> UA | aUa WT | aUa HX | aUa UA | AbdA WT | AbdA HX | AbdA UA | uAu WT | uAu HX | uAu UA |
| n | 14 | 11 | 8 | 14 | 15 | 14 | 10 | 10 | 9 | 15 | 15 | 18 |
| Mean | 78 | 66 | 79 | 75 | 62 | 73 | 95 | 83 | 79 | 50 | 38 | 48 |
| SD | 6 | 8 | 6 | 7 | 8 | 6 | 3 | 7 | 11 | 16 | 12 | 14 |
| Motif contribution* | HX to Ub<sub>x</sub> | UA to Ub<sub>x</sub> |  | HX to aUa | UA to aUa |  | HX to AbdA | UA to AbdA |  | HX to uAu | UA to uAu |  |
|  | 12 | -1 |  | 13 | 2 |  | 12 | 16 |  | 12 | 2 |  |

\*Motif contribution = Mean of WT prot activity - Mean of motif mutated prot activity

Oligos

| Construct | cDNA matrix used | primer Name | Primer sequence (5'-3') |
| --- | --- | --- | --- |
| Ubx<sup>UA</sup> | Ubx<sup>WT</sup> | Ubx5'EcoRI | CCGGAATTCATGAACCTCGTACTTTG |
|  |  | UbxUA5' | ATCAAGGCGGCGCTGCACAGGAGAAGCAGGCG |
|  |  | UbxUA3' | CTCCTGTGCAGCCGCCCTTGATCGCTGGAT |
|  |  | Ubx3'KpnI | GGGGTACCCCTACTGATCTAAGTGTC |
| Ubx<sup>OX</sup> | Ubx<sup>WT</sup> | Ubx5'EcoRI | CCGGAATTCATGAACCTCGTACTTTG |
|  |  | UbxHX5' | GCAATCACACATTGCCGCCGCGGGCTATCGCAGG |
|  |  | UbxHX3' | ATAGCCGCCGCGGCGGAATGTGTGATTGCTGGCCTGG |
|  |  | Ubx3'KpnI | GGGGTACCCCTACTGATCTAAGTGTC |

| Construct | cDNA matrix used | primer Name | Primer sequence (5'-3') |
| --- | --- | --- | --- |
| aUa | Nter | AbdA<sup>WT</sup> | Vect-a Nter Fw |
|  |  | Ubx<sup>WT</sup> | aU Nter Rv |
|  | CB | Ubx<sup>WT</sup> | aU CB Fw |
|  |  | Ubx<sup>WT</sup> | Ua CB Rv |
|  | Cter | AbdA<sup>WT</sup> | Ua Cter Fw |
|  |  | Ubx<sup>WT</sup> | u-Vect Cter Rv |

| Construct | cDNA matrix used | primer Name | Primer sequence (5'-3') |
| --- | --- | --- | --- |
| aUa<sup>OX</sup> | Nter | AbdA<sup>WT</sup> | Vect-a Nter Fw |
|  |  | Ubx<sup>OX</sup> | aU HX Nter Rv |
|  | CB | Ubx<sup>OX</sup> | aU HX CB Fw |
|  |  | Ubx<sup>OX</sup> | Ua CB Rv |
|  | Cter | AbdA<sup>WT</sup> | Ua Cter Fw |
|  |  | Ubx<sup>OX</sup> | u-Vect Cter Rv |

| Construct | cDNA matrix used | primer Name | Primer sequence (5'-3') |
| --- | --- | --- | --- |
| aUa<sup>UA</sup> | Nter | AbdA<sup>WT</sup> | Vect-a Nter Fw |
|  |  | Ubx<sup>UA</sup> | aU Nter Rv |
|  | CB | Ubx<sup>UA</sup> | aU CB Fw |
|  |  | Ubx<sup>UA</sup> | Ua UA CB Rv |
|  | Cter | AbdA<sup>WT</sup> | Ua UA Cter Fw |
|  |  | Ubx<sup>UA</sup> | u-Vect Cter Rv |

| Construct | cDNA matrix used | primer Name | Primer sequence (5'-3') |
| --- | --- | --- | --- |
| uAu | Nter | Ubx<sup>WT</sup> | Vect-u Nter Fw |
|  |  | Ubx<sup>WT</sup> | uA Nter Rv |
|  | CB | Ubx<sup>WT</sup> | uA CB Fw |
|  |  | Ubx<sup>WT</sup> | Au CB Rv |
|  | Cter | Ubx<sup>WT</sup> | Au Cter Fw |
|  |  | Ubx<sup>WT</sup> | u-Vect Cter Rv |

| Construct | cDNA matrix used | primer Name | Primer sequence (5'-3') |
| --- | --- | --- | --- |
| uAu<sup>OX</sup> | Nter | Ubx<sup>WT</sup> | Vect-u Nter Fw |
|  |  | Ubx<sup>WT</sup> | uA Nter Rv |
|  | CB | Ubx<sup>WT</sup> | uA HX CB Fw |
|  |  | Ubx<sup>WT</sup> | Au CB Rv |
|  | Cter | Ubx<sup>WT</sup> | Au Cter Fw |
|  |  | Ubx<sup>WT</sup> | u-Vect Cter Rv |

| Construct | cDNA matrix used | primer Name | Primer sequence (5'-3') |
| --- | --- | --- | --- |
| uAu<sup>UA</sup> | Nter | Ubx<sup>WT</sup> | Vect-u Nter Fw |
|  |  | Ubx<sup>WT</sup> | uA Nter Rv |
|  | CB | Ubx<sup>UA</sup> | uA CB Fw |
|  |  | Ubx<sup>UA</sup> | Au CB Rv |
|  | Cter | Ubx<sup>WT</sup> | Au Cter Fw |
|  |  | Ubx<sup>WT</sup> | u-Vect Cter Rv |

| Construct | cDNA matrix used | primer Name | Primer sequence (5'-3') |
| --- | --- | --- | --- |
| Ubx<sup>UAAbdA</sup> | Nter | Ubx<sup>WT</sup> | Vect-u Nter Fw |
|  |  | Ubx<sup>WT</sup> | UbxUAabdARv |
|  | Cter | Ubx<sup>WT</sup> | UbxUAabdAFw |
|  |  | Ubx<sup>WT</sup> | u-Vect Cter Rv |

Quantification of CRISPR AbdA::HA and Ubx::HA

| Embryo |  |  |
| --- | --- | --- |
|  | AbdA | Ubx |
| 1 | 48,28 | 73,992 |
| 2 | 63,678 | 119,954 |
| 3 | 84,799 | 84,398 |
| 4 | 79,286 | 69,162 |
| 5 | 55,915 | 75,938 |
| 6 | 34,207 | 50,064 |
| 7 | 36,363 | 77,013 |
| 8 | 51,709 | 59,876 |
| 9 | 62,245 | 28,941 |
| 10 | 83,249 | 72,216 |
| 11 | 45,718 | 46,753 |
| 12 | 81,008 | 55,829 |
